# Hepatocyte-specific miR-33 deletion attenuates NAFLD-NASH-HCC progression

**DOI:** 10.1101/2023.01.18.523503

**Authors:** Pablo Fernández-Tussy, Jonathan Sun, Magdalena P. Cardelo, Nathan L. Price, Leigh Goedeke, Chrysovalantou E. Xirouchaki, Xiaoyong Yang, Oscar Pastor-Rojo, Anton M. Bennett, Tony Tiganis, Yajaira Suárez, Carlos Fernández-Hernando

## Abstract

The complexity of the multiple mechanisms underlying non-alcoholic fatty liver disease (NAFLD) progression remains a significant challenge for the development of effective therapeutics. miRNAs have shown great promise as regulators of biological processes and as therapeutic targets for complex diseases. Here, we study the role of hepatic miR-33, an important regulator of lipid metabolism, during the progression of NAFLD. We report that miR-33 is overexpressed in hepatocytes isolated from mice with NAFLD and demonstrate that its specific suppression in hepatocytes (miR-33 *HKO*) improves multiple aspects of the disease, including insulin resistance, steatosis, and inflammation and limits the progression to non-alcoholic steatohepatitis (NASH), fibrosis and hepatocellular carcinoma (HCC). Mechanistically, we find that hepatic miR-33 deficiency reduces lipid biosynthesis and promotes mitochondrial fatty acid oxidation to reduce lipid burden in hepatocytes. Additionally, miR-33 deficiency improves mitochondrial function, reducing oxidative stress. In miR-33 deficient hepatocytes, we found an increase in AMPKα activation, which regulates several pathways resulting in the attenuation of liver disease. The reduction in lipid accumulation and liver injury resulted in decreased transcriptional activity of the YAP/TAZ pathway, which may be involved in the reduced progression to HCC in the *HKO* livers. Together, these results suggest suppressing hepatic miR-33 may be an effective therapeutic approach at different stages of NAFLD/NASH/HCC disease progression.

## INTRODUCTION

Non-alcoholic fatty liver disease (NAFLD) is the most common chronic liver disease in the world, affecting around 25% of the global population (1-8). NAFLD ranges from non-alcoholic fatty liver (NAFL) to non-alcoholic steatohepatitis (NASH) and can progress to severe fibrosis or cirrhosis and end-stage liver disease or HCC (9, 10). The rapid increase in NAFLD/NASH prevalence has paralleled the rise of obesity and diabetes, shifting NASH to the fastest growing cause of HCC in the World, especially in Western populations (3, 9-12). While the driving force of hepatic steatosis is the accumulation of fat in the liver, NAFL progression to NASH is influenced by a wide variety of factors, including genetics, inflammation, oxidative stress, mitochondrial malfunction, endoplasmic reticulum (ER) stress, lipotoxicity, insulin resistance, and gut dysbiosis (3, 13, 14).

Despite the global health and economic burden associated with NAFLD/NASH, there are still no approved therapies. Therefore, finding new potential therapeutic options is sorely needed to halt the progression of the disease and its rapid growth in the world (15, 16). Several studies have associated NAFLD with multiple metabolic maladaptations (9, 17-19). Impaired mitochondrial function is one of the most prominent metabolic alterations observed with NAFLD. Mitochondria are the most important metabolic organelles that carries out oxidative metabolism, a process that encompasses numerous pathways, including fatty acid ß-oxidation (FAO), tricarboxylic acid (TCA) cycle, electron transport chain (ETC) and adenosine triphosphate (ATP) generation. Mitochondrial dysfunction can differ depending on the stage of NAFLD, but frequently includes alterations in mitochondrial number, mtDNA, mitochondrial biogenesis, mitochondrial dynamics, and mitochondrial recycling (18, 20-26). A coordinated regulation of these processes is necessary to properly boost mitochondrial activity without detrimental effects associated with mitochondrial-derived oxidative stress and reactive oxygen species (ROS) formation. On the other hand, targeting *de* novo lipogenesis (DNL) has also arisen as a therapeutic option to temper NAFLD pathogenesis (27-30). Therefore, due to its complexity and the necessity to hit multiple pathways (15, 31), combination therapies may be the most effective approaches to treat NAFLD

MicroRNAs (miRNAs) have shown great promise as potential therapeutic targets for the treatment of metabolic disease, due to their ability to target many mRNAs and pathways simultaneously (32, 33). Previous work from our group and others identified miR-33 as an intronic miRNA hosted within the sterol regulatory element-binding protein 2 (*Srebf2*) gene (34-36). miR-33 has been shown to be an important regulator of metabolism through the regulation of mRNA transcripts involved in a wide variety of metabolic processes, including lipid and glucose metabolism (34-41). Notably, miR-33 coordinates the expression of genes associated with mitochondrial function and homeostasis (38, 42) and increased miR-33 levels in the liver (43) and serum (44) have been associated with NAFLD in humans.

Here, we elucidate a major role of hepatocyte miR-33 in regulating obesity-driven NAFL-NASH-HCC progression. Genetic ablation of hepatic miR-33 (*HKO)* improves metabolic function in the liver, enhancing glucose tolerance and insulin sensitivity and attenuating dyslipidemia, fatty liver, and NASH. In the long term, these improvements contribute to reduced liver injury and HCC development. Mechanistically, we found that hepatocyte-specific knockout of miR-33 increases mitochondrial oxidative metabolism and alters mitochondrial dynamics, which correlates with increased activation of the AMPKα signaling pathway. miR-33 regulation of AMPKα contributes to the regulation of a subset of downstream targets, including Caspase6 and TAZ, which have been recently implicated in NASH progression (45-50). Overall, this work indicates that the specific deletion of miR-33 in hepatocytes is sufficient to regulate several pathways altered throughout the development of NAFL/NASH/HCC, impeding the progression of the disease.

## RESULTS

### Loss of hepatic miR-33 improves glucose tolerance, insulin sensitivity and dyslipidemia during obesity-driven NAFL

In order to study the specific role of hepatic miR-33 in NAFL and its progression to NASH and HCC, we used the previously generated conditional miR-33 knock-out murine model (*miR-33*^*loxP/loxP*^) bred with an Albumin-Cre to induce miR-33 deletion specifically in hepatocytes (*HKO*) (51). WT and *HKO* littermates were then fed a choline-deficient, high-fat diet (CD-HFD) for 3, 6 and 15 months to induce simple steatosis/NAFL, steatohepatitis/NASH and HCC, respectively **(Supplemental Figure 1)**, as previously described (52).

To investigate the impact of hepatic miR-33 on steatosis (non-alcoholic fatty liver, NAFL), we first analyzed systemic metabolism and liver function in WT and *HKO* mice after 3 months on a CD-HFD **(Fig. 1A)**. qPCR analysis of freshly isolated hepatocytes confirmed miR-33 deletion in *HKO* mice, while revealing increased miR-33 levels in diet-induced NAFL in control mice **(Fig. 1B)**. These findings correlate with recent studies showing enhanced *SREBP2* (the host gene of miR-33a) transcriptional activation in humans and other mouse models of NAFL (53). We further confirmed this observation by measuring *SREBP2* and *SREBP1* levels in core liver biopsies from obese non-steatotic (BMI; 36-61, NAS = 0), obese steatotic (BMI; 36-61, NAS = 1-2) and obese NASH (BMI; 36-61; NAS>5, fibrosis score = 1-2). The results shown that *SREBP1* and *SREBP2* expression were markedly elevated in obese steatotic and obese NASH subjects compared to obese healthy individuals **(Supplemental Figure 1)**. As fatty liver and CD-HFD-induced NAFLD models have been associated with other metabolic dysfunctions, including obesity, dyslipidemia and insulin resistance, we next sought to determine whether miR-33 deficiency in hepatocytes influenced obesity-driven NAFLD progression (52). We observed that *HKO* mice gain less weight compared to WT mice **(Fig. 1C)**, which was accompanied by decreased body fat accumulation **(Fig. 1D)**. Circulating lipids, including total cholesterol and HDL-cholesterol were also moderately reduced in *HKO* mice while no changes were observed in circulating triglycerides (TAGs) **(Fig. 1E-H)**. Finally, we assessed the regulation of glucose homeostasis and insulin sensitivity in WT and *HKO* mice by performing glucose and insulin tolerance tests (GTT and ITT). We found that *HKO* mice showed improved glucose metabolism after 3 months on a CD-HFD **(Fig. 1I, J)**. These results agree with our previous study showing improved systemic metabolism in *HKO* mice and reinforces the metabolic benefit of depleting miR-33 in hepatocytes, independent of the underlying dietary factors driving fatty liver progression (51).

**Figure 1.**
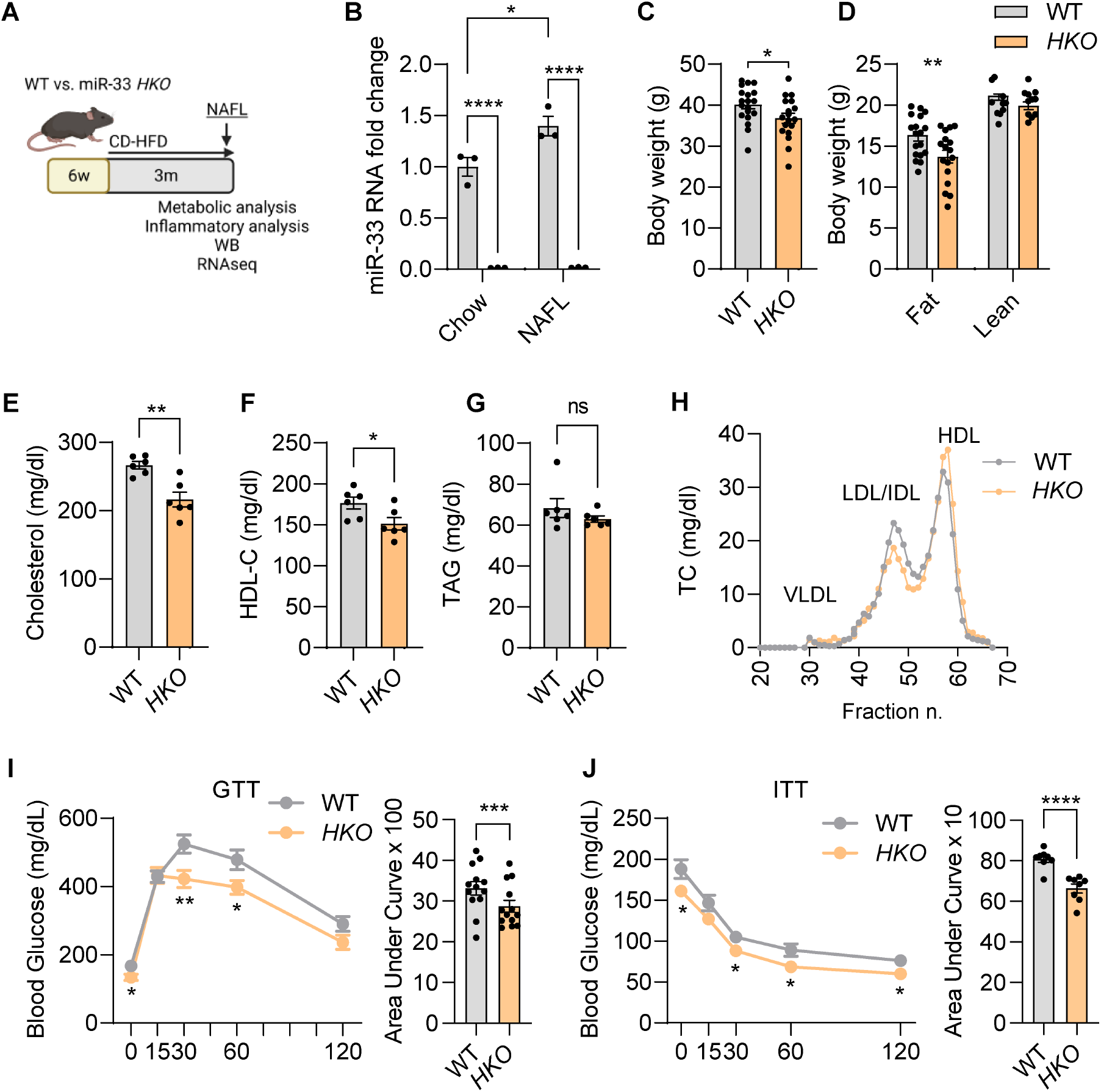
miR-33 deficiency in hepatocytes improves systemic metabolism in NAFL. **(A)** Schematic representation of the experimental design to analyze steatosis/NAFL in WT and hepatocyte specific miR-33 knockout (*HKO)* mice fed with CD-HFD for 3 months. **(B)** qPCR analysis of miR-33 expression in WT and *HKO* hepatocytes fed a control and CD-HFD for 3 months. **(C, D)** Body weight **(C)** and body composition **(D)** analysis WT and *HKO* mice. **(E-G)** Levels of total cholesterol **(E)**, HDL-C **(F)**, and TAGs **(G)** in plasma of WT and *HKO* mice. **(H)** Cholesterol content of FPLC-fractionated lipoproteins from pooled plasma of 6 WT and 6 *HKO* mice. **(I, J)** GTT **(I)** and ITT **(J)** in WT and *HKO* mice with areas under the curve. Data represent the mean ± SEM (*P ≤ 0.05, **P ≤ 0.01, ***P ≤ 0.001, ****P ≤ 0.0001 compared with WT animals, 2-way ANOVA followed by Tukey’s multiple comparison **(B)** and unpaired Student’s *t* test for 2 group comparisons).

### Genetic ablation of miR-33 in hepatocytes reduces liver steatosis by enhancing FAO and decreasing fatty acid synthesis

Hepatic energy imbalance with concurrent fat accumulation initiates NAFLD (15). Excess hepatic lipid accumulation results from the dysregulation of one or more pathways leading to an imbalance between lipid uptake, synthesis and oxidation (9). Thus, we aimed to determine whether *HKO* mice are protected against NAFLD. Our results showed a marked reduction in steatosis after feeding mice a CD-HFD for 3 months. Liver/body weight ratio and TAG content were reduced in *HKO* mice compared to WT mice, which was further confirmed by liver H&E and Oil Red O staining **(Fig. 2 A-C)**. miR-33 is an important post-transcriptional regulator of numerous genes that participate in FAO (37, 54), thus we first sought to determine if the regulation of FAO was occurring in our model of steatosis. *Ex vivo* analysis of the rate of [^14^C]-palmitate oxidation showed increased liver FAO in *HKO* mice **(Fig. 2D)**. We further characterized the contribution of miR-33 to mitochondrial metabolism by measuring the respiratory capacity of freshly isolated hepatocytes from CD-HFD fed WT and *HKO* mice using a Seahorse Bioanalyzer. This analysis further confirmed the increase in mitochondrial respiration in hepatocytes lacking miR-33 **(Fig. 2E)**. Mechanistically, we observed that carnitine O-octanoyltransferase (CROT) and the mitochondrial fatty acid (FA) transporter, CPT1a, both *bona fide* molecular targets of miR-33 and key molecules that participate in FAO, were significantly upregulated in *HKO* livers **(Fig. 2F)**.

**Figure 2.**
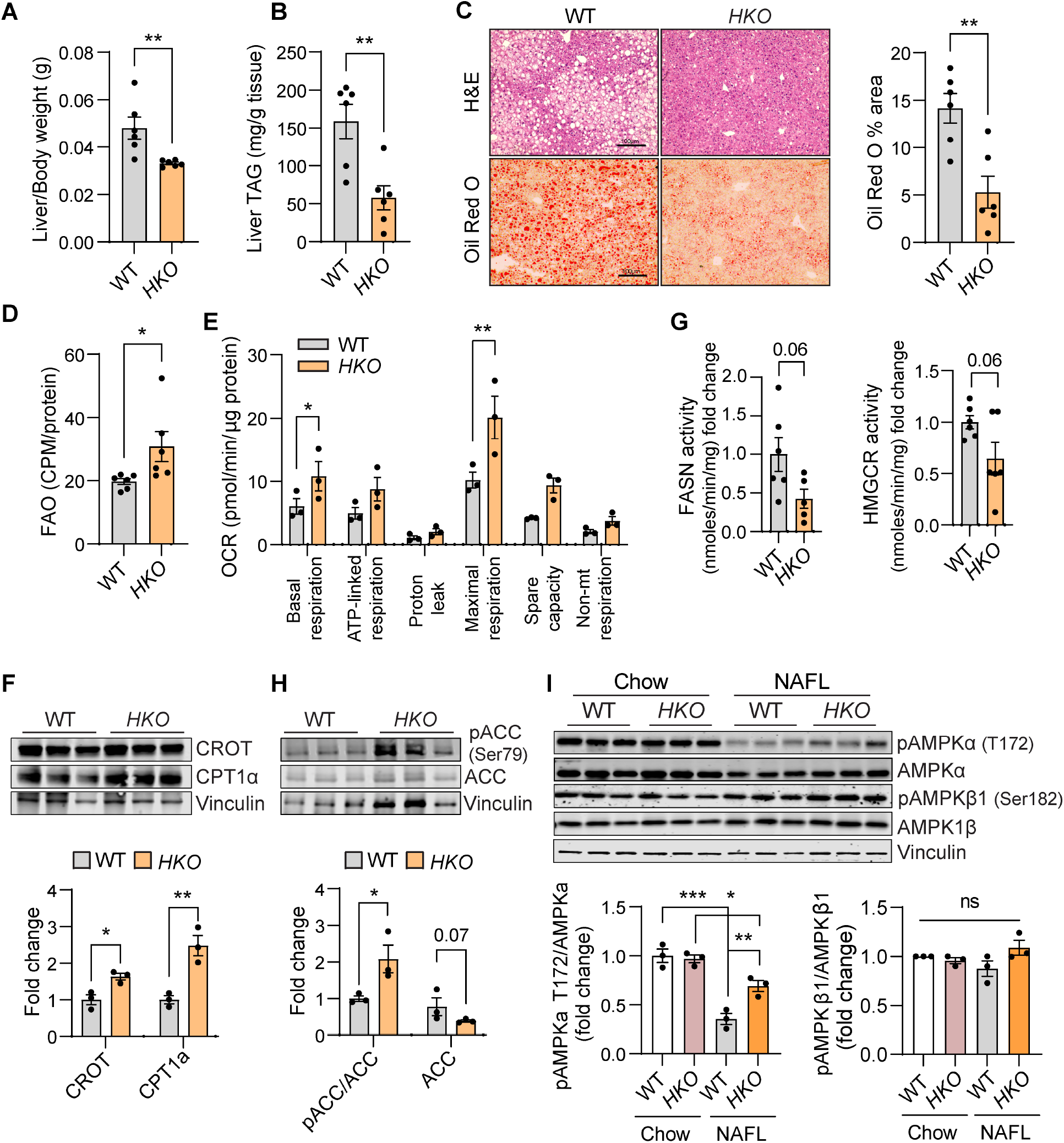
miR-33 deficiency in hepatocytes reduces liver steatosis and regulates metabolic pathways. **(A, B)** Liver weight **(A)** and liver TAG **(B)** in WT and hepatocyte specific miR-33 knockout (*HKO*) mice fed with CD-HFD for 3 months. **(C)** Representative images of H&E and Oil Red O (ORO)-stained livers from WT and *HKO* mice and quantification of ORO staining. **(D)** *Ex vivo* Analysis of FAO in WT and *HKO* livers. **(E)** Mitochondrial respiratory analysis inferred from OCR measurements of primary mouse hepatocytes isolated from WT and miR-33 *HKO* livers. **(G)** Enzymatic activity of FASN and HMGCR in WT and *HKO* liver microsomes. **(F**,**H**,**I)** Western blot and densitometric analysis of CROT, CPT1α, pACC (Ser79), total ACC, pAMPKα (T172), total AMPKα, pAMPKß (Ser182), AMPKß and housekeeping standard VINCULIN in WT and *HKO* livers. Data represent the mean ± SEM (*P ≤ 0.05, **P ≤ 0.01, ***P ≤ 0.001 compared with WT animals, unpaired Student’s *t* test for 2 group comparisons and 2-way ANOVA followed by Tukey’s multiple comparison **(I)**).

Next, we aimed to determine whether hepatocyte miR-33 deficiency influenced *DNL* during NAFLD progression. To this end, we assessed the activities of fatty acid synthase (FASN) (the enzyme involved in the synthesis of FAs from acetyl-CoA and malonyl-CoA) and HMG-CoA reductase (HMGCR) (the rate-limiting enzyme for cholesterol synthesis) in freshly isolated liver homogenates from WT and *HKO* mice. The results showed a strong trend toward decreased activity of both enzymes in *HKO* livers **(Fig. 2G)**. Consistent with this, we observed that *HKO* livers had increased Ser79 phosphorylation of Acetyl-CoA carboxylase (ACC) by AMPKα that inactivates ACC, the rate limiting enzyme for *DNL* **(Fig. 2H)**. The increased hepatic FAO and suppression of DNL observed in *HKO* mice correlated with a significant increase in AMPKα levels and activation (phosphorylation) **(Fig. 2I)**. In contrast, the expression and phosphorylation of AMPKβ was not altered, suggesting the specificity of this pathway is related to AMPKα in our model **(Fig. 2I)**.

Given the profound metabolic alterations observed in miR-33 *HKO* livers, we next assessed global transcriptomic changes by RNA-seq analysis in the livers of WT and *HKO* mice aiming to identify specific genes or upstream regulators involved in these functions. We found 1082 differentially expressed genes (DEGs) (421 up-regulated and 661 down-regulated in *HKO, Padj*. <0.05), indicating the broad effect that miR-33 deficiency has in the liver during steatosis initiation **(Fig. 3A)**. Interestingly, specific transcriptome analysis for genes involved in metabolic functions and pathways altered in obesity-driven NAFLD, revealed that gene signatures associated with FA uptake, FA synthesis and cholesterol homeostasis were altered in *HKO* livers **(Fig. 3B, C)**. Among these, *Abca1* and *Cyp7a1* upregulation in *HKO* livers were of interest, given their known role in cholesterol and bile acid metabolism and their direct regulation by miR-33. Finally, we further interrogated our RNA-seq data for changes in well-known specific processes associated with NAFLD progression, including inflammatory, profibrogenic and CYP450 associated functions (55, 56). We observed downregulation of genes associated with inflammation and fibrogenesis in *HKO* livers, while repression of CYP expression was prevented **(Fig. 3D)**. Overall, our analysis suggests miR-33 *HKO* mice are protected from NAFLD progression through the global regulation of metabolic function, including increased FAO and mitochondrial function and decreased DNL and FA uptake.

**Figure 3.**
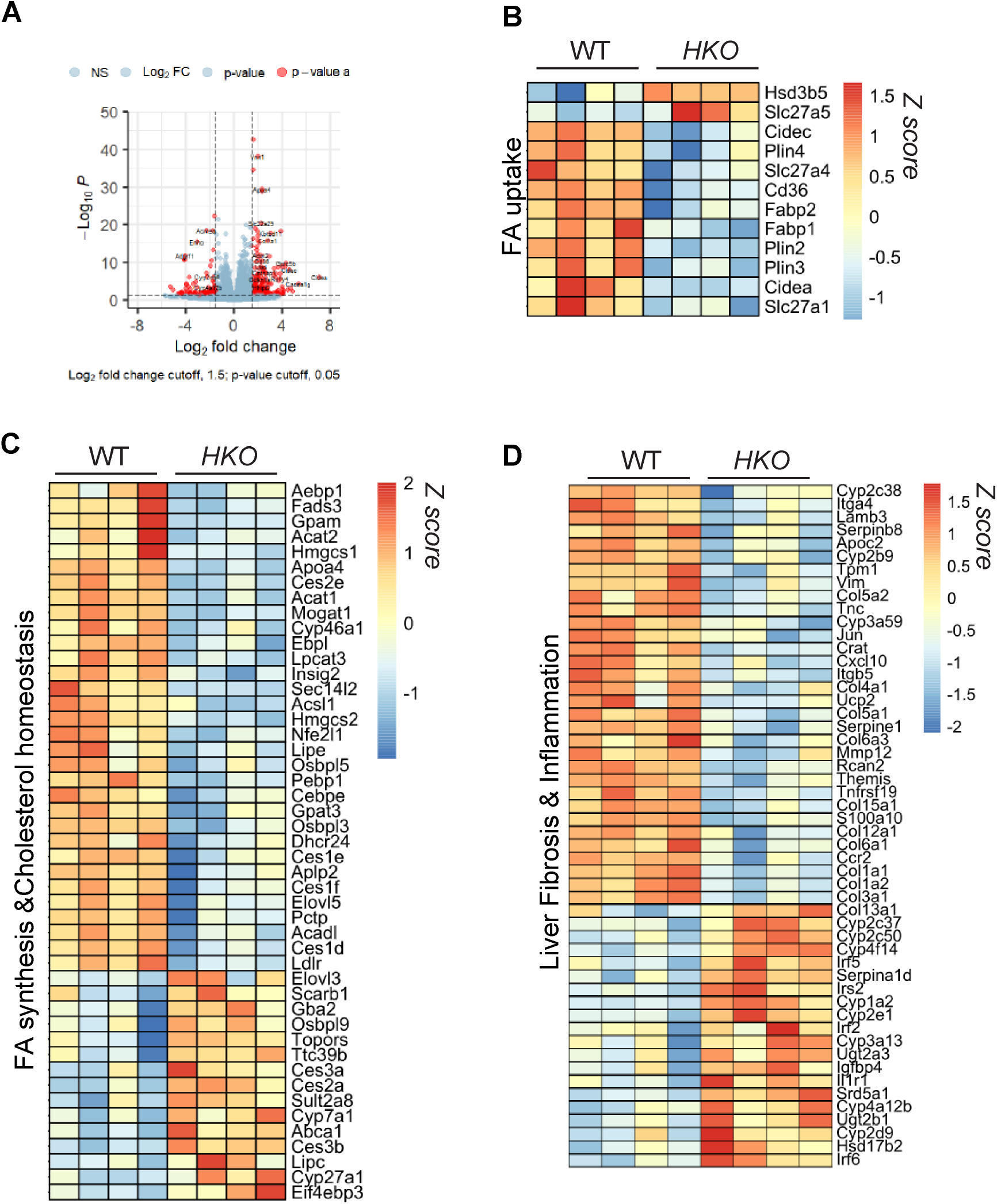
RNA-seq in NAFL livers reveals global changes in gene expression regulated by miR-33. **(A)** Volcano plot showing significant DEGs (*Padj* < 0.05, change > Log_2_ fold 1.5) in hepatocyte specific miR-33 knockout (*HKO*) vs. WT livers from mice fed with CD-HFD for 3 months. **(B-D)** Heatmaps of pathways relevant to NAFLD progression in livers from WT and *HKO* mice. Cutoff values were settled as Fold change > Log_2_1.5 and *Padj* < 0.05.

### miR-33 deficiency in hepatocytes sustains improved systemic metabolism in diet-induced NASH

The adverse outcomes associated with NASH and the ensuing fibrosis include the progression to cirrhosis and end stage liver disease or HCC (9). Halting this progression is still an unmet challenge for the development of NASH therapies. Thus, we aimed to explore whether miR-33 deficiency in hepatocytes was sufficient to improve NASH. To this end, WT and *HKO* mice were fed a CD-HFD for 6 months **(Fig. 4A and Supplemental Fig. 1)**. While miR-33 levels where still upregulated in NASH-derived hepatocytes compared to littermate controls **(Fig. 4B)**, the mild effect on body weight was no longer apparent, although body fat was reduced **(Fig. 4C, D)**. Similarly, a reduction in cholesterol levels was also observed in these mice, but to a lesser extent when compared to three months of HFD-CD feeding **(Fig. 4E-H)**. Notably, *HKO* mice showed improved glucose tolerance and insulin sensitivity **(Fig. 4I, J)**.

**Figure 4.**
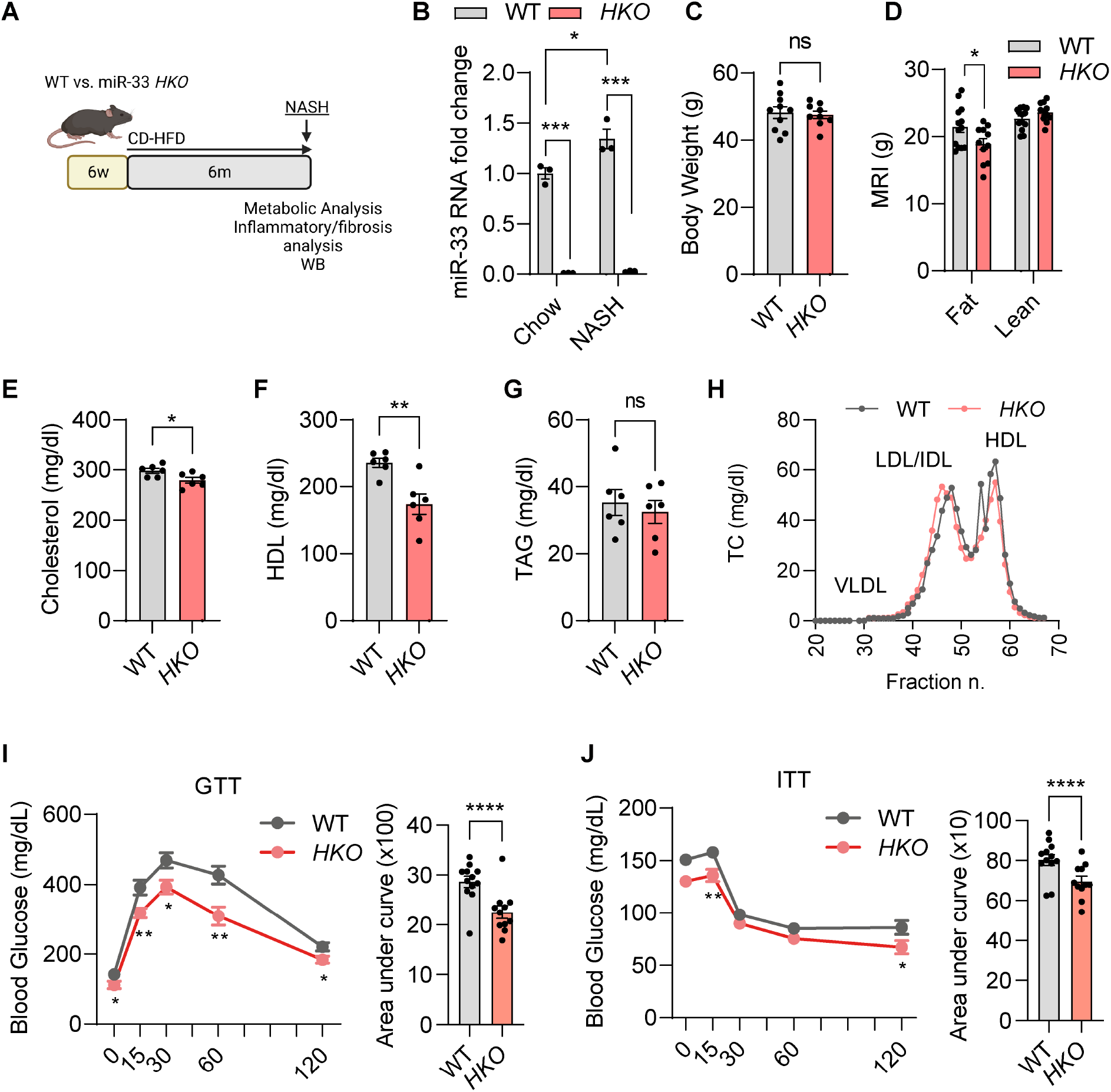
Systemic metabolism changes in miR-33 *HKO* mice at NASH stage. **(A)** Schematic representation of the experimental design to analyze steatosis/NAFL in WT and hepatocyte specific miR-33 knockout (*HKO)* mice fed with a CD-HFD for 6 months. **(B)** qPCR analysis of miR-33 expression in WT and *HKO* hepatocytes fed a control or CD-HFD for 6 months. **(C**,**D)** Body weight **(C)** and body composition **(D)** analysis in WT and *HKO* mice. **(E-G)** Levels of total cholesterol **(E)**, HDL-C **(F)**, and TAGs **(G)** in plasma of WT and *HKO* mice. **(H)** Cholesterol content of FPLC-fractionated lipoproteins from pooled plasma of 6 WT and 6 *HKO* mice. **(I, J)** GTT **(I)** and ITT **(J)** in WT and *HKO* mice with areas under the curve. Data represent the mean ± SEM (*P ≤ 0.05, **P ≤ 0.01, ***P ≤ 0.001, ****P ≤ 0.0001 compared with WT animals, 2-way ANOVA followed by Tukey’s multiple comparison **(B)** and unpaired Student’s *t* test for 2 group comparisons).

### miR-33 *HKO* mice are protected from diet-induced NASH and fibrosis

Besides regulation of systemic metabolism, we examined how miR-33 specifically affects the liver during NASH. Liver to body weight ratio was reduced in *HKO* mice, and a similar trend was observed in liver TAG content, counteracting the effect of the obesogenic diet **(Fig. 5A, B)**. NAFL progression to NASH, a more advanced disease stage, is characterized by a numerous factors, including macrovesicular fat accumulation, hepatocyte ballooning, inflammation and hepatocyte death, which results in liver damage and repair leading to different degrees of fibrosis (13). Immunohistochemistry analysis of H&E-stained liver sections revealed decreased macrovesicular fat content and hepatocyte ballooning in *HKO* livers **(Fig. 5C**), which was further confirmed by lower liver fat content and fibrosis measured by Oil Red O (ORO) and Sirius Red staining, respectively **(Fig. 5D)**. Consistent with these findings, we observed a significant reduction of liver fibrosis markers including Fibronectin (FN1), Collagen type α1 (COL1a1) and total hydroxyproline content in *HKO* mice **(Fig. 5E, F)**. Attenuation of liver fibrosis in the absence of hepatic miR-33 was not accompanied by significant reduction in liver inflammation as shown by IHC staining of F4/80+ hepatic macrophages (**Fig. 5D)** and flow cytometry analysis blood and liver leukocytes, consistent with flow cytometry analysis after 3 months of CD-HFD (**Supplemental Fig. 2 and 3**). We only observed slight changes in CD4^+^ T-cells and neutrophil presence in the livers. Reduction in liver injury in mice lacking miR-33 in hepatocytes was also confirmed by measuring serum levels of alanine aminotransferase (ALT) **(Fig. 5G)**. Together, our findings suggest that miR-33 deficiency in hepatocytes protects from diet-induced liver injury and progression of the disease to a NASH and fibrotic stage.

**Figure 5.**
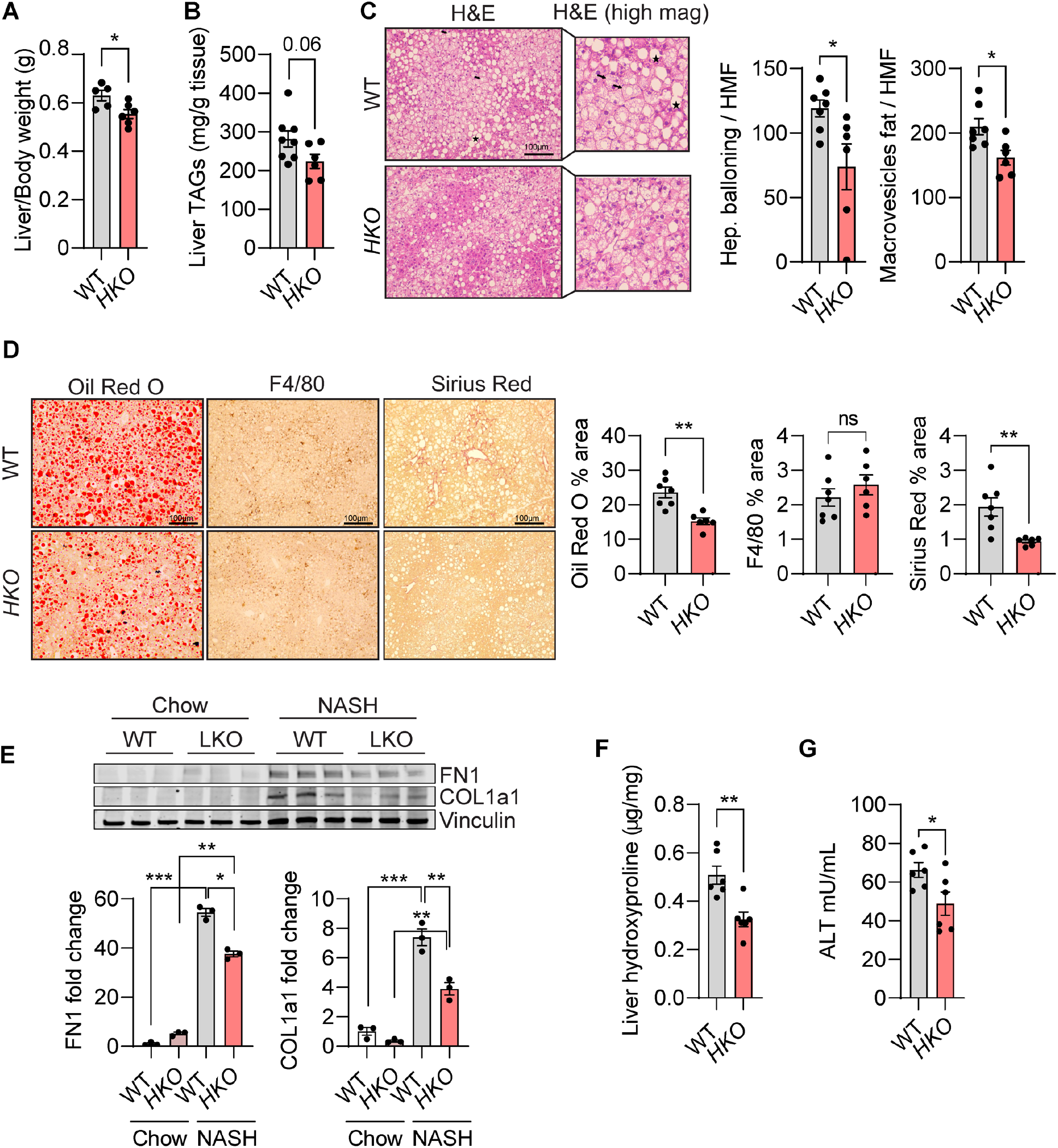
Loss of hepatic miR-33 protects from liver injury and NASH. **(A** and **B)** Liver weight **(A)** and liver TAG **(B)** in WT and hepatocyte specific miR-33 knockout *(HKO)* mice fed with CD-HFD for 6 months. **(C-D)** Representative images of H&E, Oil Red O (ORO), F4/80 and Sirius Red-stained livers from WT and *HKO* mice. Indicated quantification on the right. **(E)** Western blot and densitometric analysis of FN1, COL1a1 and housekeeping standard VINCULIN in WT and *HKO* livers fed chow or CD-HFD for 6 months (indicated as Chow and NASH, respectively in the panel). **(F)** Hydroxyproline content in NASH WT and *HKO* livers. **(G)** Serum ALT in NASH WT and *HKO* mice. Data represent the mean ± SEM (*P ≤ 0.05, **P ≤ 0.01, ***P ≤ 0.001 compared with WT animals, unpaired Student’s *t* test for 2 group comparisons and 2-way ANOVA followed by Tukey’s multiple comparison **(E)**.

We next characterized the liver metabolic adaptations of *HKO* mice in the context of NASH. Although we observed increased FAO and decreased FAs in *HKO* mice during the initial stage, it was not clear whether this improvement could be sustained over the time to contribute to improved metabolism and liver health. *Ex* vivo measurement of FAO, as well as mitochondrial respiration, established that these processes were increased in the liver and hepatocytes from *HKO* mice **(Fig. 6A, B)**. In accordance, protein levels of the miR-33 targets CROT and CPT1α were substantially increased in *HKO* livers **(Fig. 6 D)**. Similar to the early stage of disease, we also found a decrease in *DNL*, as assessed by measuring FASN/HMGCR activity and ACC protein expression **(Fig. 6C, E)**. These metabolic changes were sustained again by the upregulation of AMPKα activation in *HKO* livers **(Fig. 6F)**. Overall, this data suggests that miR-33 deficiency in hepatocytes is sufficient to increase FAO and decrease FAs, alleviating lipid overload and mitigating liver injury over a prolonged period of diet induced obesity.

**Figure 6.**
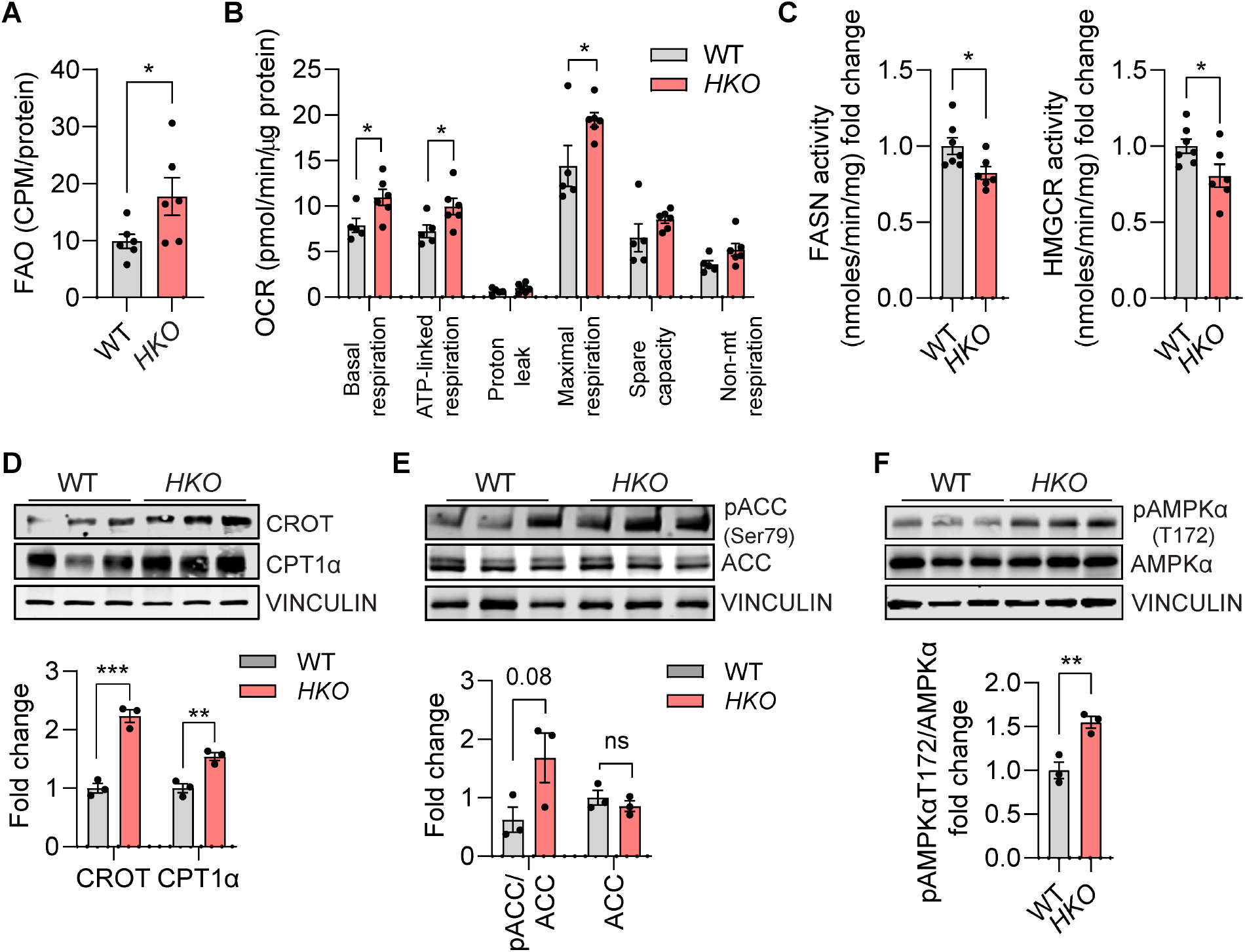
Metabolic characterization of miR-33 *HKO* mice at NASH stage. **(A)** *Ex vivo* Analysis of FAO in WT and hepatocyte specific miR-33 knockout *(HKO)* livers from mice fed with CD-HFD for 6 months. **(B)** Mitochondrial respiratory analysis inferred from OCR measurements of primary mouse hepatocytes isolated from WT and miR-33 *HKO* livers. **(C)** Enzymatic activity of FASN and HMGCR in WT and *HKO* livers. **(D-F)** Western blot and densitometric analysis of CROT, CPT1α, pACC (Ser79), total ACC, pAMPKα (T172), total AMPKα and housekeeping standard VINCULIN in WT and *HKO* livers. Data represent the mean ± SEM (*P ≤ 0.05, **P ≤ 0.01, ***P ≤ 0.001 compared with WT animals, unpaired Student’s *t* test for 2 group comparisons).

### miR-33 deficiency in hepatocytes prevents mitochondrial dysfunction associated with NAFLD/NASH progression

Mitochondrial dysfunction underlies the progression of NAFLD-NASH (18, 20-26). Previous work from our group and others identified miR-33 as a master regulator of mitochondrial function through the targeting of several genes involved in mitochondrial biogenesis, metabolism, and homeostasis (41, 42, 57). Notably, we found that miR-33 ablation in hepatocytes improves their metabolic function and mitochondrial respiratory capacity even under conditions of prolonged mitochondrial stress. We next aimed to further characterize the molecular mechanism that mediates the improvement in mitochondrial function observed in miR-33 deficient hepatocytes. We found increased mitochondrial content in hepatocytes from *HKO* NASH livers, measured by protein levels of different complexes of the electron transport chain (ETC) and assessing mitochondrial to nuclear DNA ratio (mtDNA/nDNA) **(Fig. 7A and Supplemental Fig. 4A)**. This effect was found in CD-HFD challenged mice, but not in lean mice fed a chow diet **(Supplemental 4Fig. B**,). These findings were further supported by electron microscopy analysis of hepatocytes from NASH mice, which revealed an increase in the coverage and density of mitochondrial mass, as well as mitochondrial elongation in *HKO* mice **(Fig. 7C-G)**. We also observed enhanced mitochondrial ETC activity of Complex I and Complex II **(Fig. 7B)**. The increase in mitochondrial mass found in hepatocytes from *HKO* mice correlated with elevated levels of PGC1α, a transcription factor targeted by miR-33, mainly known for its role in promoting mitochondrial biogenesis (58, 59). Moreover, its downstream target TFAM, was also upregulated in miR-33-deficient livers, suggesting increased mitochondrial biogenesis **(Fig. 7H)**. Together, these results demonstrate that absence of miR-33 in hepatocytes improves mitochondrial function increasing mitochondrial mass and ETC activity.

**Figure 7.**
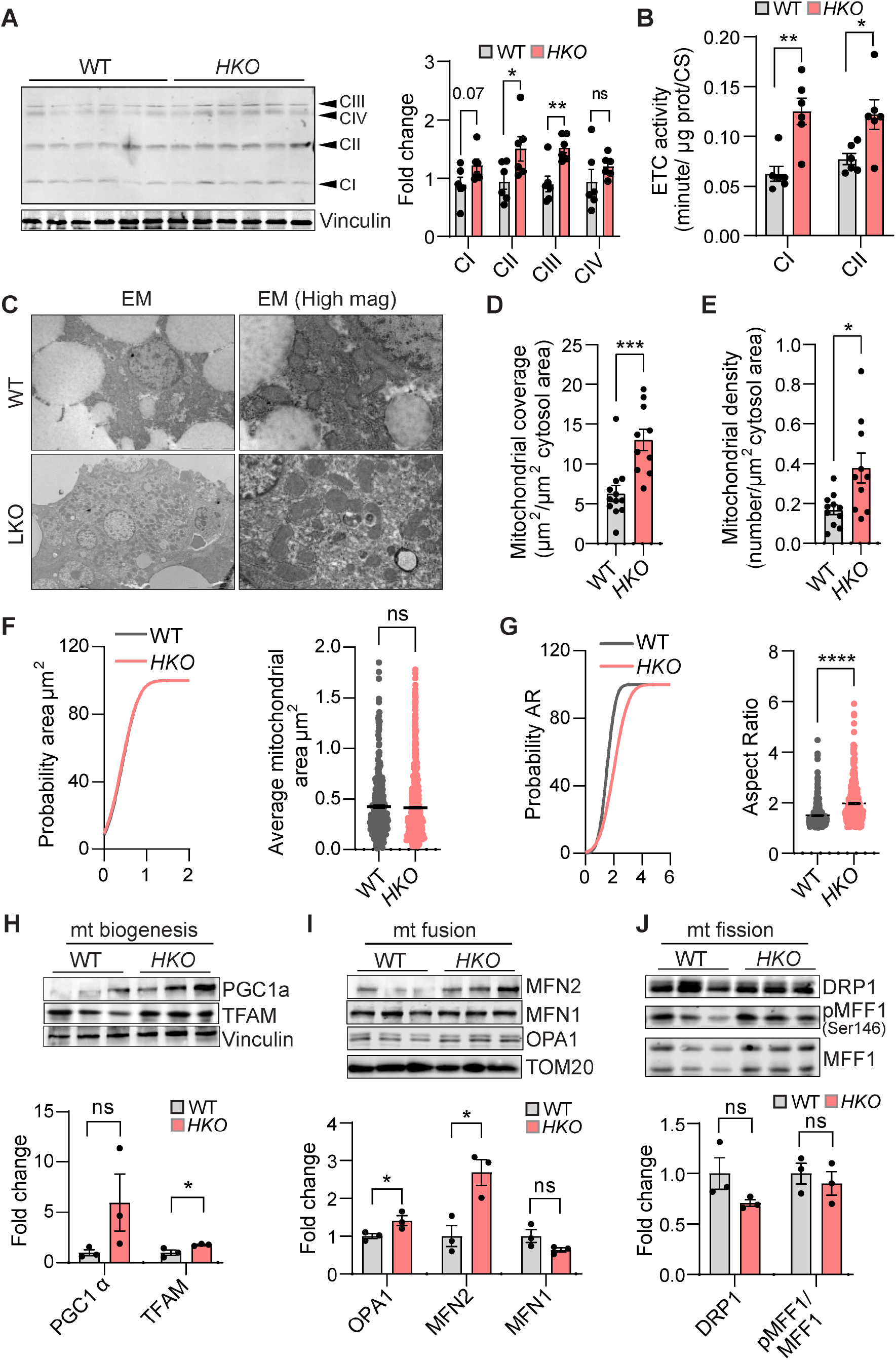
Impact of miR-33 deficiency on mitochondrial homeostasis and morphology. **(A)** Western blot and densitometric analysis of different mitochondrial subunits blotted with the Total OXPHOS Rodent WB Antibody Cocktail (ab110413) and housekeeping standard VINCULIN in WT and hepatocyte specific miR-33 knockout (*HKO*) livers from mice fed with CD-HFD for 6 months. **(B)** Activity of the ETC Complex I and Complex II in NASH livers. Enzyme activities are expressed as change in absorbance / minute / μg protein / CS activity. **(C)** Representative electron micrographs of mitochondria profiles in WT and *HKO* hepatocytes from NASH livers. **(D-G)** Mitochondrial coverage **(D)**, Mitochondrial density **(E)** and cumulative distribution and mean of mitochondrial area **(F)** and mitochondria aspect ratio **(G)** from WT and *HKO* hepatocytes. **(H-J)** Western blot and densitometric analysis of **(H)** PGC1α and TFAM; **(I)** MFN2, MFN1, OPA1; **(J)** DRP1, p-MFF1 (Ser146), MFF1 and housekeeping standard VINCULIN or TOM20 in WT and *HKO* livers. Data represent the mean ± SEM (*P ≤ 0.05, **P ≤ 0.01, ***P ≤ 0.001, ****P ≤ 0.0001 compared with WT animals, unpaired Student’s *t* test for 2 group comparisons).

Mitochondrial homeostasis is a critical checkpoint for the control of mitochondrial health and metabolism (18, 24, 60). Mitochondrial quality control mechanisms include mitochondrial biogenesis, mitochondrial dynamics to balance fusion and fission processes, and mitophagy. Dysregulations of all these processes is thought to facilitate NAFLD progression (22, 60). Thus, we sought to characterize these processes in our NASH model. Mitochondrial number and size are also controlled through the balance of mitochondrial dynamics, a process that involves fusion and fission of mitochondrial membranes and has been described in NAFLD (24, 61). We measured the levels of the most relevant proteins participating in the regulation of fusion/fission and found an increase in the fusion related proteins MFN2 and OPA1, but no relevant changes in fission proteins **(Fig. 7I, J)**. Importantly, the increase observed in MFN2 levels supports the changes in mitochondrial shape observed by EM, and correlates also with the increased respiratory capacity of these mice.

Lipid overload and excessive mitochondrial activity have been linked to mitochondrial dysfunction in NAFLD. Besides the inability to sustain metabolic needs, mitochondrial dysfunction is responsible for the production of large amounts of reactive oxygen species (ROS), which increases mitochondrial damage and eventually lead to cell death (62). Although increased mitochondrial number and activity in *HKO* mice could lead to higher ROS production and damage, changes in mitochondrial dynamics can also play a role in ROS regulation, membrane potential and other downstream processes related to mitochondrial stress (24, 60). To determine whether miR-33 levels in hepatocytes influence ROS production in obesity-driven NAFLD/NASH, we monitored ROS accumulation in mitochondria from liver sections and observed a decrease in *HKO* mice **(Fig. 8A)**. Liver lipid peroxidation measured by assessing malondialdehyde (MDA) as a readout of ROS damage also showed a similar decrease in livers from *HKO* mice **(Fig. 8B)**. No significant changes were found in the oxidized or reduced forms of glutathione, or their ratio, and levels of glutathione peroxidase 4 and peroxiredoxin were also unaffected by loss of miR-33 **(Supplemental Fig. 5A, B and)**. However, we noticed a marked increase in glutathione-reductase activity, a marker of reduced oxidative stress in *HKO* livers, suggesting that changes in the recycling rather than the synthesis of glutathione may contribute to reduced oxidative stress in these livers **(Fig. 8C)**. We also observed nuclear erythroid 2-related factor 2 (NRF2) and downstream targets such as NQO-1 and HO-1 were also increased in *HKO* mice, while Keap1, which prevents NRF2 from translocating to the nucleus was downregulated **(Fig. 8D)**. Considering the close link between mitochondrial dynamics, dysfunction, lipid overload and ER stress, we interrogated *HKO* NASH livers for changes in ER stress response. However, no significant changes were found in support of a role of miR-33 in regulating ER stress **(Supplemental Fig. 5C)**. Finally, the ultimate cellular consequence of mitochondrial dysfunction and oxidative stress, the induction of cell death, was also attenuated in the *HKO* mice under NASH conditions, as seen by caspase activity and TUNEL staining **(Fig. 8E-G and Supplemental Fig. 5D)**. Together, these findings indicate that miR-33 deficiency in hepatocytes improves mitochondrial quality control enhancing mitochondrial biogenesis and mitochondrial dynamics, to sustain high rates of oxidative metabolism without increasing mitochondrial injury and oxidative stress during lipid overload, protecting against hepatocyte cell death.

**Figure 8.**
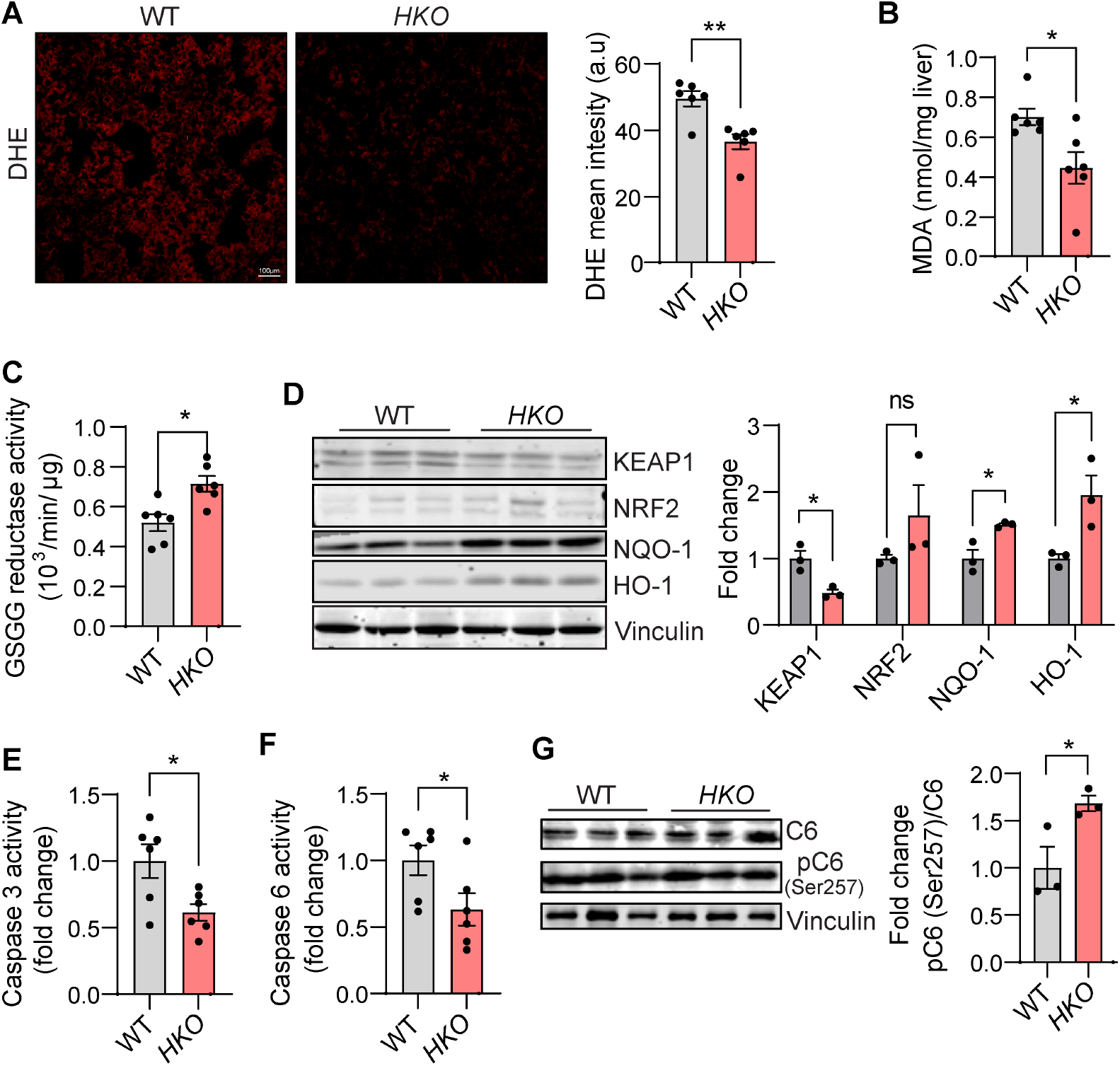
Genetic ablation of miR-33 in hepatocytes attenuates hepatic oxidative stress and cell death in NASH. **(A)** Representative DHE staining and quantification in WT and hepatocyte specific miR-33 knockout (*HKO*) livers from mice fed with CD-HFD for 6 months. **(B)** Lipid peroxidation measured by MDA assay in NASH livers. **(C)** Glutathione reductase activity measured in NASH livers. Data represented as change in absorbance / minute/ μg of protein). **(D)** Western blot and densitometric analysis of KEAP1, NRF2, NQO-1 and HO-1 and housekeeping standard VINCULIN in Wt and *HKO* livers. **(E, F)** Fold change in Caspase 3 and Caspase 6 activity in NASH livers. **(G)** Western blot analysis and densitometric analysis of p-Caspase 6 (Ser257) and Caspase 6 and housekeeping standard VINCULIN in WT and *HKO* NASH livers. Data represent the mean ± SEM (*P ≤ 0.05, **P ≤ 0.01, compared with WT animals, unpaired Student’s *t* test for 2 group comparisons).

### AMPK signaling pathway is increased in miR-33 *HKO* livers

AMPK is a master regulator of metabolism and mitochondrial homeostasis (63). Our previous results showed that AMPK activation is increased in *HKO* mice compared to WT mice in both NAFL and NASH stages, counteracting the progressive decrease reported in NAFLD (45). These results prompted us to characterize additional posttranscriptional mechanisms driving AMPK regulation in our model. Notably, we found that the activation of liver kinase B1 (LKB1), a kinase that controls AMPK activity, was enhanced as shown by the increased phosphorylation of LKB1 at serine 428 in *HKO* livers **(Fig. 9A)**. LKB1 activation is regulated by its subcellular compartment through deacetylation and phosphorylation (64-66), which correlates with increased levels of protein deacetylases of sirtuins, including SIRT1, SIRT2, SIRT3, SIRT7 and a trend toward upregulation of SIRT6 **(Fig. 9B)**. Sirtuin activity is dependent not only on expression levels but also on the availability of NAD^+^. Accordingly, we detected that total NAD, NAD^+^ and NAD^+^/NADH were increased in *HKO* livers **(Fig. 9C Supplemental Fig. 6A)**.

**Figure 9.**
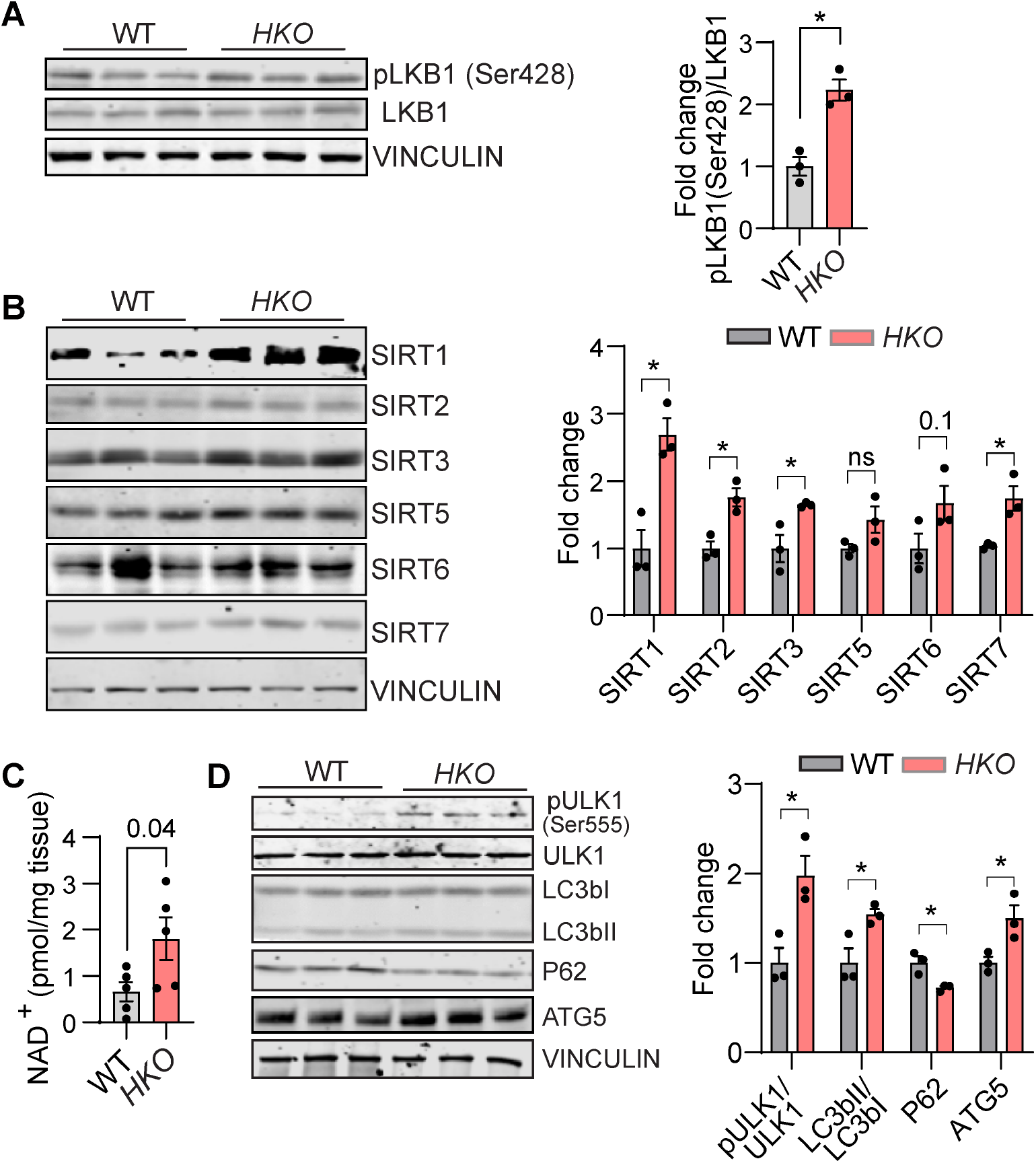
Upstream and downstream AMPKα signaling pathway is altered in hepatocyte deficient miR-33 livers. **(A, B)** Western blot analysis and densitometric analysis of **(A)** p-LKB1 (Ser428) (long and short exposure), LKB1; **(B)** SIRT1, SIRT2, SIRT3, SIRT5, SIRT6, SIRT 7 and housekeeping standard VINCULIN in WT and hepatocyte specific miR-33 knockout (*HKO*) livers from mice fed with CD-HFD for 6 months. **(C)** NAD^+^ levels in WT and *HKO* NASH livers represented as pmol/mg of tissue. **(D)** Western blot analysis and densitometric analysis of p-ULK1 (Ser555), ULK1, LC3bI/II, P62/SQSTM1, ATG5 and housekeeping standard VINCULIN in WT and *HKO* NASH livers. Data represent the mean ± SEM (*P ≤ 0.05, **P ≤ 0.01, compared with WT animals, unpaired Student’s *t* test for 2 group comparisons).

These results point towards the increased activation of upstream regulators of AMPKα. As previously shown in Figure 6, we found increased FAO and decreased FAs, with increased AMPKα/ACC phosphorylation indicating a broad rewiring of metabolism mediated by AMPK signaling in *HKO* livers. Similarly, we aimed to characterize if other metabolic pathways regulated by AMPK were altered in mice lacking miR-33 in hepatocytes. Consistent with our observations on the effects of AMPK, we found that ULK1 phosphorylation, a downstream target of AMPKα was also increased in *HKO* livers, which along with the increased expression of LC3bII and ATG5 and decreased expression of P62, indicates an increased autophagy flux in *HKO* livers **(Fig. 9D)**.

### miR-33 deficiency in hepatocytes reduces NAFLD progression to HCC

To analyze whether the improved metabolic function and protection against NAFLD-NASH progression attenuates the HCC incidence in *HKO* mice, we fed WT and *HKO* mice a CD-HFD for 15 months. Tumor quantification revealed a marked decrease in the tumor incidence and average number of tumors per mouse in *HKO* mice **(Fig. 10A** and **B)**, which was particularly pronounced in larger tumors (volume >20mm^3^) **(Fig. 10C)**. In agreement with reduced tumor incidence, serum levels of α-fetoprotein (AFP), a common circulating marker for HCC, were significantly reduced in *HKO* mice compared to WT mice **(Fig. 10D)**. Histological analysis of WT and *HKO* tumors also revealed a decrease in proliferative Ki67 positive cells in tumors from *HKO* mice compared to WT mice **(Fig. 10E)**. Recent studies have highlighted the role of the gene regulator TAZ in NASH worsening and progression to HCC (46, 47, 50, 67, 68). YAP/TAZ are transcriptional coactivators of the Hippo pathway that participate in the initiation and progression of different cancers (68-70). Specifically, TAZ levels in HCC have been associated with its initiation and prognosis (50, 67, 71). TAZ upregulation in NASH and HCC has been associated with both increased cholesterol levels and decreased AMPKα activity and it has been described to participate in the transcriptional regulation of several genes involved in fibrosis, proliferation, superoxide formation and regulation of metabolism. Its upregulation has been described in pre-tumor NASH stage (50, 67, 71). In accordance with these studies, we also found a marked upregulation of TAZ levels in NASH livers that was partially abrogated in mice lacking miR-33 in hepatocytes **(Fig. 10F)**. We further confirmed the activation of TAZ in NASH livers by cellular fractionation and immunoblotting for its nuclear localization **(Fig. 10G)**. In line with this, downstream TAZ target genes were decreased in *HKO* livers, further suggesting the downregulation of this pathway **(Fig. 10H)**. Based upon our results, we speculated two different mechanisms could be involved in TAZ downregulation in *HKO* livers: i) YAP/TAZ is a direct target of AMPK and phosphorylation of TAZ and its partner YAP was increased in *HKO* livers compared to WT **(Fig. 10I)**; ii) cholesterol accumulation in NASH mediates TAZ stabilization and subsequent activation and we have observed a significant decrease of both total and free cholesterol in *HKO* livers **(Fig. 10J)**, which may also contribute to its regulation. Taken together, our present results suggest miR-33 deficiency in hepatocytes improves mitochondrial metabolic function, restraining NAFLD/NASH progression and in the long-term, preventing the development of HCC **(Fig. 11)**.

**Figure 10.**
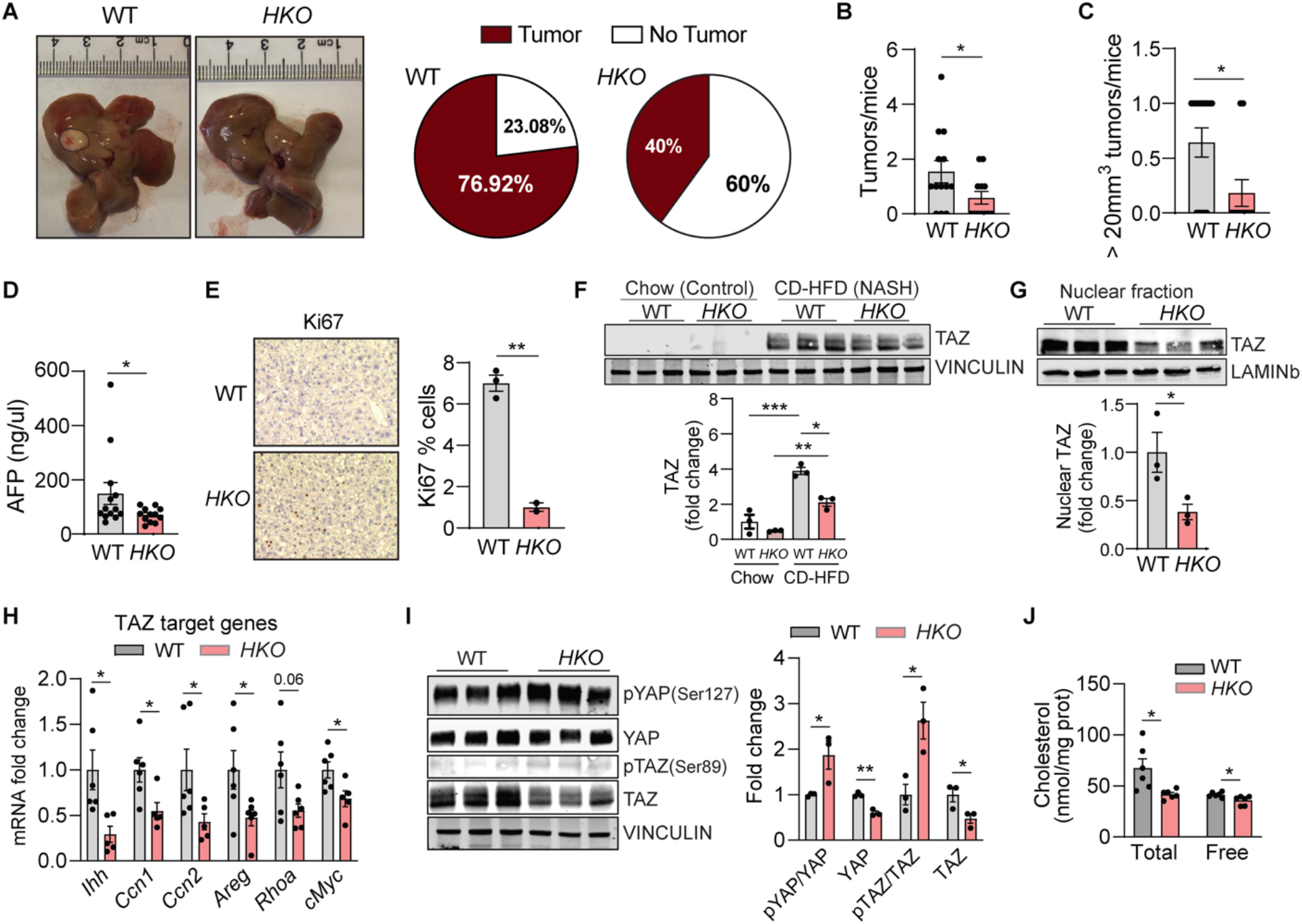
Absence of hepatocyte miR-33 decreased Hippo signaling pathway and hepatic tumor incidence. **(A)** Representative images of WT and hepatocyte specific miR-33 knockout (*HKO*) livers after 15 months of CD-HFD, dashed line used to outline tumors (right panel). Graphical representation of tumor incidence in WT and HKO mice (left panel). **(B)** Total number of tumors/mouse and **(C)** Number of tumors larger >20mm^3^/mouse found in WT and *HKO* livers after 15 months of CD-HFD. **(D)** Circulating AFP levels in WT and *HKO* mice after 15 months of CD-HFD. **(E)** Representative images of Ki67 staining in liver tumors from WT and *HKO* mice after 15 months on CD-HFD. Quantification is shown on the right panel. **(F)** Western blot and densitometric analysis of TAZ and housekeeping standard VINCULIN in WT and *HKO* livers fed chow and CD-HFD for 6 months. **(G)** Western blot and densitometric analysis of TAZ and housekeeping standard LAMINb in nuclear liver fractions from WT and *HKO* liver after 6 months of CD-HFD. **(H)** qPCR analysis of mRNA expression of TAZ-responsive genes including *Ihh, Ccn1, Ccn2, Areg, Rhoa, cMYC* in WT and *HKO* livers after 6 months on CD-HFD. **(I)** Western blot and densitometric analysis of p-YAP (Ser127), YAP, p-TAZ (ser89) TAZ, and housekeeping standard VINCULIN in WT and *HKO* livers after 6 months on CD-HFD. **(J)** Hepatic total and free cholesterol levels in WT and HKO mice fed a CD-HFD for 6 months of measured by GC-MS. Data represented as nmol of cholesterol/mg of liver protein. Data represent the mean ± SEM (*P ≤ 0.05, **P ≤ 0.01, ***P ≤ 0.001 compared with WT animals, unpaired Student’s *t* test for 2 group comparisons).

**Figure 11.**
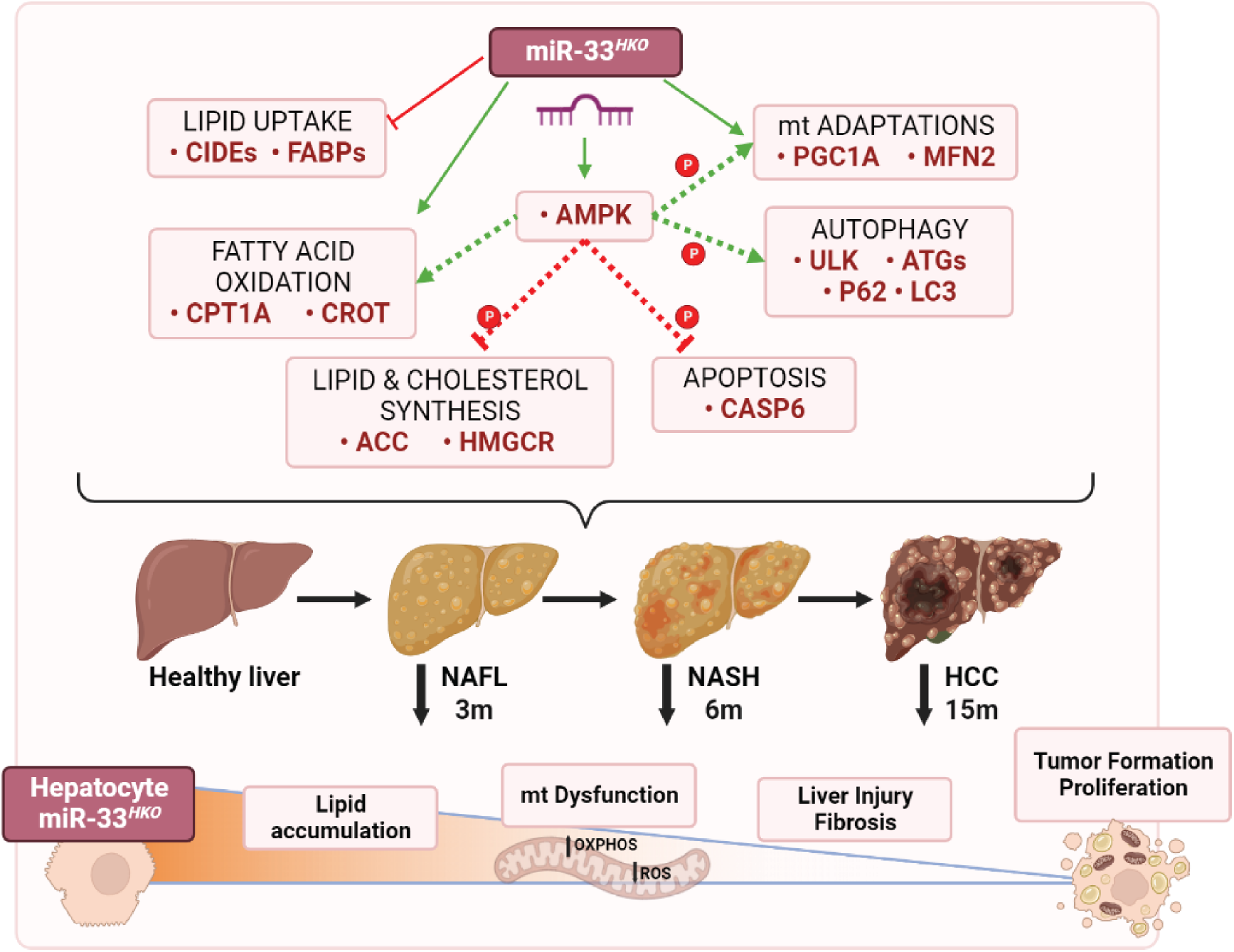
Schematical representation of miR-33 mechanism of action.

## DISSCUSION

While the rise in NAFLD and overnutrition approaches pandemic levels in Western societies, the complexity of the disease has hindered the development of viable therapeutic options for its treatment. In this study, we demonstrate that miR-33 expression is increased in hepatocytes at different stages of NAFLD and that the specific deletion of miR-33 in hepatocytes improves liver function, reducing lipid accumulation in the liver and the progression of the disease. Work from numerous groups has demonstrated the role of miR-33 in metabolism, including cholesterol biosynthesis and efflux, triglyceride metabolism, autophagy, glucose and insulin homeostasis and mitochondrial function (34-42). Those data demonstrated that the inhibition or deletion of miR-33 was sufficient to reduce the development of atherosclerotic plaques in mice and non-human primates (38, 39, 72-77). However, due to the promiscuous nature of miRNAs, whole-body deficiency of miR-33 was associated with obesity, dyslipidemia and insulin resistance (78, 79). These detrimental effects held up further studies investigating the role of miR-33 in other metabolic diseases, such as NAFLD. Recently, new strategies to overcome potential undesired effects of miRNA therapies have been investigated, shedding light on the cell-specific functions of miR-33 and its therapeutic value. Some of these studies, have demonstrated the efficiency of delivering miR-33 inhibitors inside pH low insertion peptides (pHLIP) to the kidney and atherosclerotic lesions (80, 81). Moreover, in a recent study, using a different strategy, our group also demonstrated the safety and efficiency of specifically removing miR-33 from hepatocytes to improve cholesterol and FA metabolism, highlighting the role of hepatic miR-33 in liver metabolism and fibrosis (51). This previous study not only showed that miR-33 suppression in hepatocytes was not responsible for the adverse metabolic effects observed in whole-body deficient mice, but that liver specific loss of miR-33 improved whole body metabolism under hyperlipidemic conditions (51). Taking advantage of this model, in this work, we have focused on the metabolic benefits associated with miR-33 deficiency in hepatocytes, during NAFLD/NASH/HCC development to interrogate the long-term alterations in liver function under this chronic inflammatory disease.

The initial characterization of miR-33 *HKO* in this CD-HFD model showed a clear impact on the regulation of cholesterol and glucose metabolism, with almost no effect on body weight, consistent with previous results found for *HKO* mice on other diets (51). The decrease in steatosis was related to the regulation of several pathways involved in lipid accumulation in the liver, including FA uptake, DNL and FAO, as evidenced by *ex* vivo metabolic assays, RNA-sequencing and protein level analysis performed in *HKO* livers. Further assessment confirmed that *HKO* mice showed improved liver function regarding glucose and lipid metabolism in advanced stages of NAFLD, protecting from disease progression in the long term. Our data suggest that these improvements in liver metabolism that alleviate lipid buildup are primarily responsible for ameliorating liver injury and progression of the disease, as no other predicted miR-33 targets associated with fibrosis were found to be dysregulated in our model.

One of the most common features in NAFLD and obesity is the inability to sustain mitochondrial adaptation to nutrient status (18, 20-26, 61). Mitochondria are highly dynamic organelles with the ability to undergo functional and structural changes in response to environment and energy requirements (60). However, in NAFLD, as with other metabolic diseases, high FAO rates to counteract lipid accumulation often results in rapid increases of oxidative stress and ER stress, resulting in mitochondrial injury, defective oxidative phosphorylation, and impaired energy production (18, 26, 61, 82). Here, we describe that not only FAO but also oxidative phosphorylation was in increased miR-33 *HKO* hepatocytes. However, we did not find increase in oxidative stress was detected in these mice. Further interrogation of hepatic mitochondria from WT and *HKO* mice fed a CD-HFD for 6 months showed that miR-33 deficiency is associated with dramatic changes in mitochondrial quantity and morphology, suggesting a broader role of miR-33 in metabolism beyond regulation of cholesterol and FAO. The observed changes in mitochondria suggest miR-33 *HKO* mice have increased mitochondrial biogenesis and mitochondrial dynamics, mechanisms directed by PGC1α and MFN2 among other markers. This mitochondrial phenotype is associated with increased oxidative capacity along with reduced ROS production and inflammation, resulting in protection from liver injury (24-26, 60). The combination of the different approaches used here to study mitochondrial turnover and dynamics suggests both a positive regulation of mitochondrial biogenesis and mitochondrial fusion; however, as these processes are usually connected to each other, we cannot discard the dominance of one over the other. Thus, the observed increase in mtDNA and ETC protein complexes could be a consequence of increased membrane fusion and recycling of mitochondrial fragments, rather than exclusively a consequence of increased mitochondrial biogenesis. Moreover, as mitochondria are highly dynamic organelles that respond to the cellular energy needs, we cannot discount the possibility that the different regulatory processes could occur at the same time in these livers depending on the specific requirements. Although our data points to a phenotype with a greater number and more elongated mitochondria to support FAO and ETC activity, a role for miR-33 in mitochondrial regulation through fission or mitophagy cannot be ruled-out under these circumstances, such as a situation of increased mitochondrial damage, as these processes are necessary for the recycling of mitochondrial fragments that cannot be recycled through fusion (60).

Biochemical and metabolic analyses revealed AMPKα activation as a central node in the protection from NAFLD progression in *HKO* mice. Activation of AMPKα correlated with downstream pathways, including DNL, FAO, mitochondrial function, autophagy, caspase activation and YAP/TAZ transcriptional activity (45, 63, 83, 84). YAP/TAZ activation in hepatocytes plays a central role in liver fibrosis and transition to hepatocellular carcinoma and is also known to be regulated by lipid accumulation and cholesterol levels (47). Thus, given the multiple adaptations regulated in *HKO* mice, the decrease observed in YAP/TAZ activation could be a consequence of AMPKα phosphorylation or an indirect consequence of the decreased cholesterol accumulated in these livers. The exact mechanism by which YAP/TAZ is regulated in our model remains to be further studied to know the role of AMPKα on YAP/TAZ activation in hepatocytes.

This study will contribute to a better understanding of the mechanisms involved in NAFLD/NASH progression and how therapeutic interventions could be applied to its treatment. The role of enhanced FAO as beneficial or detrimental in NAFLD/NASH has been highly discussed in the past, our results bring new insights into the beneficial role of FAO in the disease. (15, 25, 31, 85). These findings indicate that the regulation of hepatocyte metabolism by miR-33 is involved in the progression of NAFLD/NASH, as well as NAFLD/NASH-derived HCC and, that direct targeting of miR-33 in hepatocytes protects from the progression of this disease. This may be particularly relevant for the use of approaches such as N-acetylgalactosamine–conjugated antisense oligonucleotides, which have been demonstrated to be effective for targeted delivery of inhibitors to the liver (86). Regarding human pathology, it is important to note, that while mice have only the miR-33a isoform of miR-33, humans express both miR-33a and miR-33b isoforms. While miR-33a is encoded within the *SREBF2* gene, miR-33b is encoded within the *SREBF1* gene, which is regulated by different mechanisms (87, 88). Moreover, the regulation of both *SREBF1* and *SREBF2* transcriptional activity has recently been observed in both mice and human NAFLD (53), suggesting that in human pathology, the miR-33b isoform may also be contributing to the development of the disease, further escalating the therapeutic potential of targeting hepatic miR-33 in human pathology.

## METHODS

### Animals

miR-33-knock-out mice (*miR-33*^*loxP/loxP*^) were generated as previously described by our laboratory with the assistance of Cyagen Biosciences (51). To generate hepatocyte specific miR-33-knock-out mice, *miR-33*^*loxP/loxP*^ mice were bred with transgenic mice expressing *Cre* recombinase under the control of a hepatocyte-specific promoter: Albumin promoter (JAX stock 003574). To produce diet-induced liver disease, mice were fed a standard chow diet until 8 weeks of age, then chow diet was replaced by modified choline-deficient high fat diet, containing 45% of fat and no choline added (CD-HFD) (D05010402, Research Diets, USA). Mice were maintained with CD-HFD feeding for 3, 6 or 15 months to induce simple steatosis (Non-Alcoholic Fatty Liver, NAFL), steatohepatitis (NASH) or hepatocellular carcinoma (HCC) at respective time points, according to previous descriptions of the model (52). Body weight was measured throughout diet feeding studies, and analysis of body composition was performed by Echo MRI (Echo Medical System). Mice used in all experiments were sex- and age-matched and kept in individually ventilated cages in a pathogen-free facility. All mice were fasted for 6h at end time point experiments. All the described experiments were approved by the institutional animal care use committee of Yale University School of Medicine.

### Human liver biopsies

The use of human tissue was approved by the Monash University Human Research Ethics Committee (CF12/2339-2012001246; CF15/3041-2015001282). All subjects gave their written consent before participating in this study. Liver core biopsies were from obese men and women undergoing bariatric surgery have been described previously (89) and were processed for RNA isolation. Gender differences analyses were not performed due to the low frequency of suitable donors.

### Lipoprotein profile and circulating lipid measurement

Mice were fasted for 6h before blood samples were collected cardiac puncture, and plasma was separated by centrifugation. HDL-C was isolated by precipitation of non-HDL cholesterol and both HDL-C fractions and total plasma were stored at –80 °C. Total plasma cholesterol and triglycerides were measured using kits according to the manufacturer’s instructions (Wako Pure Chemicals, Japan). The lipid distribution in plasma lipoprotein fractions was assessed by fast performance liquid chromatography (FPLC) gel filtration with 2 Superose 6 HR 10/30 columns (Pharmacia Biotech, Sweden).

### Glucose and Insulin tolerance test

GTTs were performed after overnight fasting (16h) by intraperitoneal (IP) injection of glucose at a dose of 1.5g/kg. Blood glucose was measured at 0-, 15-, 30-, 60- and 120-minutes post injection. ITTs were performed following 6h fasting by IP injection of 1.5U/kg of insulin. Blood glucose was measured at 0-, 15-, 30-, 60- and 120-minutes post injection.

### Liver Triglyceride and cholesterol measurements

For determining total TAGs content in the liver, TAGs were extracted using a solvent chloroform/methanol (2:1). TAG level in the liver was determined by using a commercially available assay kit (Sekisui Chemical Co., USA) according to the manufacturer’s instructions. Determination of liver total and free cholesterol was determined gas-chromatography coupled with mass spectrometry (GC-MS). Briefly, liver tissue extracts equivalent to 1 mg/mg.Prot. were mixed with 1.5 ml of methanol/chloroform (Cl3CH) mixture (2:1, v/v) in presence of μ25 L of aqueous KOH 50% (w/v). Cholestanol (5α-cholestan-3β-ol, Sigma-Aldrich, USA) was added on every sample as internal standard. After incubation for 1h at 90ºC, the saponified lipid extract containing the total cholesterol was extracted with 1 mL of Cl3CH and 2 mL of water, the lower phase recovered and dried over nitrogen current. Cholesterol was analyzed by GC-MS as described previously (90). The same extraction protocol, without the addition of KOH, was done to extract free cholesterol from liver tissues.

### Fatty acid oxidation (FAO)

Ex vivo FAO was analyzed using [^14^C] palmitate, as previously described (91). Briefly, livers were isolated from WT and *HKO* and homogenized in five volumes of chilled STE buffer (10 mM Tris-HCl, 0.25M sucrose, and 1 mM EDTA and pH 7.4). Homogenate was centrifuged, and the pellet was incubated with reaction mixture (0.5 mmol/L palmitate conjugated to 7% BSA/[^14^C]-palmitate at 0.4μCi/ml) for 30 minutes. After this incubation, the resuspended pellet containing the reaction mixture was transferred to an Eppendorf tube, the cap of which housed a Whatman filter paper disk that had been presoaked with 1 mol/L sodium hydroxide. The ^14^C trapped in the reaction mixture media was then released by acidification of media using 1 mol/L perchloric acid and gentle agitation of the tubes at 37C for 1h. Radioactivity that had become absorbed onto the disk was then quantified by liquid scintillation counting in a ß-counter.

### MitoStress test

Real-time measurements of oxygen consumption rate (OCR) were measured as previously described using a Seahorse XF24 Extracellular Flux Analyzer (Seahorse Biosciences, USA), (92). Briefly, WT and *HKO* primary hepatocytes from CD-HFD fed mice at the indicated time points were isolated by the Yale Liver Center by standard liver perfusion and collagenase digestion followed by centrifugation of the cell suspension at 60g for 4 minutes to pellet hepatocytes. Hepatocytes were then cultured in collagen type I coated XF24 cell culture microplates (Seahorse Bioscience) at 1.5 ×10^4^ cells per well and incubated 4-6 h at 37 °C. After that, the cells were washed once in 1x PBS and media was changed to low-glucose assay media for overnight incubation at 37 °C. The next morning hepatocytes were washed twice with 1 ml XF Assay Media (DMEM base containing 1 mM pyruvate, 2 mM glutamine and 5.5 mM glucose, pH 7.4) and incubated at 37 °C for 1 h in a non-CO2 incubator. Cells were then assayed on a Seahorse XFe24 Analyzer following a 12-min equilibration period. Respiration rates were measured using an instrument protocol of 3-min mix, 2-min wait, and 3-min measure. The following inhibitors were used at the indicated concentration: oligomycin (1 μM); carbonyl cyanide 4-trifluoromethoxyphenylhydrazone (FCCP) (1 μM); and rotenone (0.5μM)/antimycin (0.5μM). Flux rates were normalized to total protein content following cell lysis at the end of the assay.

### Liver Flow Cytometry analysis

A small piece (250-300 μg) of PBS perfused liver was resected and placed in 2ml cold PBS, then clopped into smaller pieces by mechanical disruption. The homogenate was then transfer into a gentleMACS C Tube and further dissociated with a gentleMACS Dissociator (Program liver 1, x2). Homogenate was digested with liver digestion buffer (5 ml HBSS w/o Ca^2+^/Mg^2+^, 200U/ml Collagenase IV (Worthington, USA) (37 °C, 30mins, ∼60-80 rpm). After digestion, homogenate was filtered through 70 μm filter, washed with 10 ml blocking buffer (HBSS w/o Ca^2+^/Mg^2+^, 2% FBS, 5mM EDTA) and centrifuged at 500g, 5mins, RT. Pellet was resuspended in 10 ml 20% Percoll and centrifuged 1300g, 30 mins, RT. 13. Pellet was resuspended in 1mL of HBSS blocking solution and transferred to a 1.5 ml Eppendorf tube, then centrifuged at 500g, 5 mins, RT. Final pellet was resuspended in 200 μl of ACK (155 mM ammonium chloride, 10 mM potassium bicarbonate, and 0.01 mM EDTA, pH 7.4) and then stained with a mizture of antibodies. B cells were identified as APC-Cy7 B220 (Biolegend, USA); T cells were identified as CD4^hi^ or CD8^hi^ with the following antibodies: BUV395 CD90.2 - 565257 (BD, USA), BV711 CD4 - 100447 (Biolegend), BV605 CD8a - 100744 (Biolegend) and activation was determined according to CD62L/CD44 status with PE-Cy7 CD62L - 25-0621-82 (eBioscience, USA), BUV737 CD44 - 612799 (BD), BV605 CD8a - 100744 (Biolegend); macrophages were identified as FITC F480 - 157310 (Biolegend); neutrophils were identified as CD11b^hi^Ly6G^hi^ with Pacific Blue CD11b - 101224 (Biolegend), APC Ly6G - 127614 (Biolegend). All antibodies were used at 1:300 dilutions.

### Blood Flow Cytometry analysis

Blood was collected by heart puncture. For FACS analysis, erythrocytes were lysed with ACK lysis buffer (155 mM ammonium chloride, 10 mM potassium bicarbonate, and 0.01 mM EDTA, pH 7.4). White blood cells were resuspended in 3% fetal bovine serum (FBS) in PBS, blocked with 2 μg mL−1 of FcgRII/III, then stained with a mixture of antibodies. Monocytes were identified as CD115^hi^ and subsets as Ly6-C^hi^ and Ly6-C^lo^; neutrophils were identified as CD11b^hi^Ly6G^hi^; B cells were identified as B220^hi^; T cells were identified as CD4^hi^ or CD8^hi^. All antibodies were used at 1:300 dilutions.

### RNA-seq

Total RNA from livers of control and HKO mice was extracted and purified using a RNA isolation Kit (Qiagen) followed by DNAse treatment to remove genomic contamination using RNA MinElute Cleanup (Qiagen). The purity and integrity of total RNA sample was verified using the Agilent Bioanalyzer (Agilent Technologies, Santa Clara, CA). rRNA was depleted from RNA samples using Ribo-Zero rRNA Removal Kit (Illumina, USA). RNA libraries were performed TrueSeq Small RNA Library preparation (Illumina) and were sequenced for 45 cycles on Illumina HiSeq 2000 platform (1 × 75bp read length). The reads obtained from the sequencer are trimmed for quality using inhouse developed scripts. The trimmed reads were aligned to the reference genome using TopHat2. The transcript abundances and differences were calculated using cuffdiff. The results were plotted using R and cummeRbund using in-house developed scripts. RNA-sequencing data have been deposited in the Gene Expression Omnibus database (GSE220093).

### Fatty acid and Cholesterol synthesis

FASN activity was determined in the liver, as described previously with some modifications (93). Briefly, liver from WT and *HKO* mice were homogenized in tissue homogenization buffer (0.1 M Tris, 0.1 M KCl, 350 μM EDTA, and 1 M sucrose, pH 7.5) containing protease inhibitor cocktail (Roche, USA). The supernatant was collected by centrifuging liver homogenates at 9,400 g for 10 minutes at 4°C. For determining FASN activity, Liver homogenate was added to NADPH activity buffer (0.1 M potassium phosphate buffer, pH 7.5 containing 1 mM DTT, 25 μM acetyl-CoA, and 150 μM NADPH). Malonyl-CoA (50 μM) was added to the assay buffer to initiate the reaction. The decrease in the absorbance was followed at 340 nm for 30 mins at an interval of 1 min using a spectrophotometer set in the kinetic mode under constant temperature (37°C). HMGCR Activity Assay The HMG-CoA reductase activity assay was determined according to the protocol described previously (94), with slight modifications. In Brief, a microsomal fraction from cell lysates and the liver homogenate was obtained via ultracentrifugation (100000 x g for 60 min). The reaction buffer (0.16 M potassium phosphate, 0.2 M KCl, 400 μM EDTA, and 0.01 M dithiothreitol) containing 100 μM NADPH and microsomal protein (200 μg/mL) was prewarmed at 37 ºC for 10 min before the reaction. The reaction was initiated by adding 50 μM substrate (HMG-CoA) to the reaction buffer. The decrease in the absorbance at 340 nm was followed for 30 min with an interval of 1 min.

### Electron Microscopy and Mitochondrial Analysis

Sample preparation and imaging was performed by the Center for Cellular and Molecular Imaging (CCMI) Electron Microscopy Core Facility at Yale University. Briefly, Liver pieces were fixed with 2.5% glutaraldehyde and 2% PFA in 0.1 M sodium cacodylate (pH 7.4) for 2 h, RT. Cells were postfixed in 1% OsO_4_ in the same buffer for 1 h, then stained en bloc with 2% aqueous uranyl acetate for 30 min, dehydrated in a graded series of ethanol to 100%, and embedded in Poly/bed 812 for 24 h. Thin sections (60 nm) were cut with a Leica ultramicrotome and poststained with uranyl acetate and lead citrate. Digital images were taken using a Morada charge-coupled device camera fitted with iTEM imaging software (Olympus). Mitochondria analysis was performed as described previously described (95). Mitochondria cross-sectional area and mitochondria aspect ratio (major axis divided by minor axis, minimum value is 1.0) were calculated as a measurement of mitochondria size and shape, respectively. Probability plots were utilized to estimate changes in mitochondria size and shape, and statistical differences were tested using Kolmogorov–Smirnov test. Mitochondria density was estimated by dividing the number of mitochondria profiles by the cytosolic area. Mitochondria coverage was estimated by dividing the total area of mitochondria by the cytosolic area.

### DHE

Dihydroexiethidium staining was performed in 8μm sections from OCT-embedded livers. Sections were incubated with MnTBAP (150μM, 1h, RT), stained with DHE (Sigma-Aldrich) (5μM, 30 min, 37 ºC) and mounted with VECTASHIELD(R) Antifade Mounting Medium (Vector Laboratories, USA). Stained area percentage was calculated using ImageJ software.

### ETC activity complex I and Complex II

Frozen liver tissues were homogenized with 20mM KP buffer (pH 7.4) with a glass homogenizer and centrifuged at 800g, 10 mins. Enzyme activities were measured at 30 ºC, monitoring the reaction for at least 2 h and normalizing the changes in absorbance to CS activity and protein concentration. Complex I activity was measured by tracking the oxidation of NADH at 340 nm in 20mM KP buffer (pH 8).0) with 200 μM NADH, 1mM NaN_3_, 0.1% BSA-EDTA and 100 μM ubiquinone-1 with and without rotenone (5 μM), to calculate the rotenone-sensitive rate of NADH oxidation. Complex II was assayed following the reduction of 2,6-dichlorophenolindophenol (DCPIP) at 600 nm. Reaction was set in 50 mM Tris-KP (pH 7.0), with 1.5 mM KCN, 100 μm DCPIP and 32 mM succinate. Citrate synthase (CS) activity was measured at 420 nm in 75 mM Tris-HCl (pH 8.0) buffer with 100 μM DTNM, 350 μg/ml acetyl-CoA, 0.5 mM oxalacetate and 0.1% Triton X-100).

### Glutathione Reductase Activity

The reductase activity of glutathione was calculated in liver homogenates as the reduction of GSGG observed in the presence of NADPH. Briefly, livers were homogenized in assay buffer (0.2 M KPH, pH 7.0, 2 mM EDTA). Assay was performed to measure changes in NADPH absorbance at 340 nm. Reaction buffer was prepared as follows: 100 μl of assay buffer were added with 10 μl GSSG (20mM) and 10 μl NADPH (2 mM) and brought to a final volume of 200 μl with H_2_O. Sample was added to reaction solution and absorbance at 340 nm was monitored for at least 30 mins. Changes in absorbance were normalized to protein concentration in liver lysates.

### NAD/NADH

NAD/NADH were measured in liver samples with the NAD/NADH Assay Kit ab65348 (Abcam, UK) following manufacturer’s recommendations.

### Caspase-3 and Caspase-6

Caspase-3 activity was assayed as previously described (96). Briefly, livers protein lysates were incubated with fluorescent substrate Ac-DEVD-AMC Caspase-3 Fluorogenic Substrate (BD Biosciences) at 37°C and the reaction was monitored for at least 4 h to track changes in fluorescence (λ excitation of 390 nm and λ emission of 510 nm). For Caspase-6 activity assay, similar procedure was followed for colorimetric detection with Caspase-6 Colorimetric Assay Kit K115 (BioVision, USA), monitoring absorbance at 405 nm.

### Statistical analysis

All data are expressed as mean ± SEM unless indicated. Statistical differences were measured using unpaired two-sided Student’s *t* test. Normality was checked using the Kolmogorov–Smirnov test. A value of *P* ≤ 0.05 was considered statistically significant. Data analysis was performed using GraphPad Prism Software Version 9.0 (GraphPad).

### Study approval

Animal experiments were conducted under the ethical guidelines of and protocols were approved by the IACUC at Yale University School of Medicine (animal protocol 2019-11577).

## AUTHOR CONTRIBUTION

PFT, YS and CFH designed the research. PFT, JS, MPC, NLP, LG, CEX, OPR performed research and analyzed data. XY, AMB, TT analyzed data and edited manuscript. PFT and CFH wrote the manuscript.

## ACKNOWLEDGMENTS

This work was supported by grants from the National Institutes of Health (R35HL135820 to CF-H; and R35HL155988 to YS and 1K01DK120794 to NLP and R00HL150234 to LG), the American Heart Association (20TPA35490416 to CF-H and 874771 to PF-T), Programa Postdoctoral de Perfecionamiento de Personal del Gobierno Vasco (Spain) (to PF-T.).

