## Supplemental for "Hepatocyte-specific miR-33 deletion attenuates NAFLD-NASH-HCC progression"

#### SUPPLEMENTAL METHODS

**RNA isolation and Real-Time qPCR.** Total RNA, including miRNAs, was isolated from liver tissue or hepatocytes using the miRNeasy miRNA Isolation Kit according to the manufacturer's instructions and reverse-transcribed into cDNA with iscript cDNA systhesis kit (Bio-Rad, USA). qPCR was performed with SsoFast EvaGreen Supermix (Bio-Rad) using the same thermal profile conditions for all primers sets: 40 amplification cycles (95 °C for 5 s and 60 °C for 10 s). All samples were analyzed in duplicate and normalized to 18S. To analyze miR-33, RNAs were reverse-transcribed with the miRCURY LNA RT Kit (Qiagen, Germany) and quantified with the miRCURY LNA miRNA PCR assays (Qiagen). miR-33 levels were normalized to the levels of SNORD68 RNA.

**Liver Histology.** Mouse livers were perfused with PBS and fixed in 4% paraformaldehyde (PFA) overnight. Then livers were transferred to 70% for immunohistochemistry (IHC) analysis and submitted for staining through the Yale Pathology Tissue Services core, with H&E, Sirius Red, F4/80, TUNEL and Ki67 to analyze liver morphology, fibrosis, inflammation, cell death and proliferation. Additionally, an independent piece of liver was transferred from PFA to 30% sucrose for lipid staining with Oil Red O. For each staining, pictures were taken with an EVOS microscope. Hepatocyte ballooning was quantified as relative number of ballooned hepatocytes per frame. Abundance of macrovesiclular fat, Sirius Red, F4/80 and Oil Red O staining were quantified by number or area percentage with ImageJ software. TUNEL and Ki67 were quantified as number of positive nuclei per total number of nuclei.

**Western Blot analysis.** Tissues were homogenized by manual disruption and the Bullet Blender Homogenizer in ice-cold buffer containing 50mMTris·HCl, pH 7.5, 0.1% sodium dodecyl sulfate (SDS), 0.1% deoxycholic acid, 0.1 mM EDTA, 0.1 mM EGTA, 1% Nonidet P-40, 5.3 mM NaF, 1.5 mM NaP, 1 mM orthovanadate, 1 mg/mL protease inhibitor mixture (Roche), and 0.2mg/mL 4-(2-aminoethyl) benzenesulfonyl fluoride hydrochloride (Roche). Lysates were sonicated and rotated at 4 °C for 1 h before the insoluble material was removed by centrifugation at 12,000 Å~ g for 10 min. After normalizing for equal protein concentration, cell lysates were resuspended in SDS sample buffer before separation by SDS polyacrylamide gel electrophoresis. Following transfer of the proteins onto nitrocellulose membranes, the membranes were probed with the following antibodies: CROT (Novus no. 3144; 1:1,000), CPT1A (Abnova H00001374-P01; 1:1,000), ACC (Ser79) (CST #11818; 1:1000) , ACC (CST #3676; 1:1000), AMPK (T712) (CST #2535; 1:1000), AMPKα (CST #5831; 1:1000), AMPKβ1(Ser182) (CST #4186; 1:1000), AMPKβ1 (CST #4150; 1:1000), FN1 ( Sigma F3648; 1:1000), COL1a1 (Novus

Biologicals NB600; 1:500), Total OXPHOS cocktail (Abcam ab110413; 1:2000), PGC1$\alpha$ (Abcam ab54481; 1:1000), TFAM (Abcam ab252432; 1:1000), MFN1 (Abcam ab126575; 1:1000), MFN2 (Abcam ab56889; 1:1000), OPA1 (CST #80471; 1:1000), TOM20 (CST #42406; 1:1000), DRP1 (CST #8570; 1:1000), MFF1 (Ser146) (CST #49281; 1:1000), MFF1 (CST #84580; 1:1000), KEAP1 (CST #8047; 1:1000), NRF2 (CST #12721; 1:500), NQO-1 (CST #62262; 1:1000), HO-1 (CST #43966; 1:1000), C6 (Ser257), C6 (CST #9762; 1:1000), LKB1 (Ser428) (CST #3482; 1:1000), LKB (CST #13031; 1:1000), SIRT1 (CST #9475; 1:1000), SIRT2 (CST #12650; 1:1000), SIRT3 (CST #5490; 1:1000), SIRT5 (CST #8782; 1:1000), SIRT6 (CST #12486; 1:1000), SIRT7 (CST #5360; 1:1000), ULK1 (Ser555) (CST #5869; 1:500), ULK1 (Novus Biologicals NBP2-24738SS; 1:1000), LC3bI/II (CST #12741; 1:1000), SQSTM1/P62 (CST #39749; 1:1000),, ATG5 (CST #12994; 1:1000), YAP/TAZ, YAP (Ser127) (CST #13008; 1:1000), TAZ(Ser89) (CST #59971; 1:1000), GAPDH (Abcam ab8245; 1:5000), LAMINb (Abcam ab19109; 1:5000), VINCULIN (Sigma V9131; 1:5000). For preparation of cytosol and nuclear extracts from liver tissue the Nuclear Extract Kit (ab219177, Abcam, UK) was used following manufacturer's recommendation. Subsequent WB analysis were performed regularly.

**ALT Measurements.** ALT activity was determined in serum with the ALT Activity Assay Kit (Sigma-Aldrich MAK052) following the manufacturer's recommendations.

**Liver Hydroxyproline assay.** Hydroxyproline content in the liver was measured with the following protocol. Liver tissue (20-40mg) was homogenized in 6N HCl (1:20 w:v) and incubated 16-24h at 110 ºC. On the next day, livers were cooled down and filtrated with a 0.22 $\mu$m syringe filter. Homogenates were then neutralized with 2.2% NaOH in Citrate-Acetate-Buffer at a 1:10 ratio. Reaction was set as follows: 500 $\mu$l of samples was incubated with 250 $\mu$l of chloramine-T (20 min, RT), then 250 $\mu$l of PCA were added (20 min, RT), and 250 $\mu$l of Dimethylbenzaldehyde were added to the mixture and incubated (20 min, 60 ºC). Absorbance was measured at 565 nm.

**Lipid Peroxidation.** Liver lipid peroxidation was measured with the Lipid Peroxidation (MDA) assay kit MAK085 (Sigma-Aldrich), following manufacturer's recommendations for colorimetric detection at 532 nm.

**AFP Measurements.** AFP levels in serum were determined with the AFP kit - MAFP00 (R&D, USA) following manufacturer's recommendations.

#### SUPPLEMENTAL FIGURE LEGENDS

**Supplemental Figure 1. CD-HFD model in WT and *HKO* mice. (A)** Schematic representation of the experimental design to analyze whole NAFLD-NASH-HCC progression from steatosis/NAFL to NASH to HCC in WT and hepatocyte specific miR-33 knockout (*HKO*) mice fed with CD-HFD for 3-,6- and 15-months.

**Supplemental Figure 2. Srebp1 and Srebp2 expression is elevated in obese subjects with NAFLD and NASH.** qRT-PCR analysis of *SREBP1* and *SREBP2* mRNA levels in healthy obese without steatosis (n=7; NAS = 0), NAFL (n=7; NAS = 1-2) or NASH and fibrosis (n=7; NAS> 5, fibrosis score = 1-2).

**Quantitative real-time PCR (qPCR) analysis of *DUSP1* in human liver core biopsies from obese patients without steatosis (n=7; NAS = 0), NAFL (n=8; NAS = 1-2) or NASH and fibrosis (n=8; NAS> 5, fibrosis score = 1-2).** Data represent the mean ± SEM. *P ≤ 0.05 was determined by one-way ANAVO

**Supplemental Figure 3. Analysis of inflammatory markers in NAFL.** Flow cytometry analysis of blood **(A)** and liver **(B)** leukocytes from WT and hepatocyte specific miR-33 knockout (HKO) mice at 3 months of CD-HFD feeding. Data are expressed as percentages of live cells. Data represent the mean ± SEM (*P ≤ 0.05 compared with WT animals, unpaired Student's *t* test).

**Supplemental Figure 4. Analysis of inflammatory markers in NASH.** Flow cytometry analysis of blood **(A)** and liver **(B)** leukocytes from WT and HKO mice at 6 months of CD-HFD feeding. Data are expressed as percentages of live cells. Data represent the mean ± SEM (*P ≤ 0.05 compared with WT animals, unpaired Student's *t* test).

**Supplemental Figure 5. Analysis of mitochondrial content in WT and *HKO* livers.** Mitochondrial DNA to nuclear DNA ratio in WT and hepatocyte specific miR-33 knockout (*HKO*) mice at 6 months of CD-HFD feeding **(A)** and in lean chow-diet-fed mice **(B)**. Data represent the mean ± SEM (*P ≤ 0.05 compared with WT animals, unpaired Student's *t* test).

**Supplemental Figure 6. Analysis of oxidative stress, ER-stress and apoptosis in NASH. (A)** GSH and GSGG levels and ratio in livers of WT and hepatocyte specific miR-33 knockout (HKO) mice at 6 months of CD-HFD feeding. **(B)** Western blot and densitometric analysis of GPX4, PRDX and housekeeping VINCULIN in NASH livers from WT and HKO mice. **(C)** Western blot and densitometric analysis of ATF4, BIP, IRE1, PERK, XBP1s, and housekeeping VINCULIN in NASH livers from WT and HKO mice. **(D)** Representative images and quantification of TUNEL-stained NASH livers. Data represent the mean ± SEM (**P ≤ 0.01 compared with WT animals, unpaired Student's *t* test).

**Supplemental Figure 7. NADH/$NAD^+$ regulation in *HKO* livers. (A)** NADH and $NAD^+$ levels in WT and hepatocyte specific miR-33 knockout (*HKO*) livers at 6 months of CD-HFD feeding. Data represent the mean ± SEM (*P ≤ 0.05 compared with WT animals, unpaired Student's *t* test).

**A**

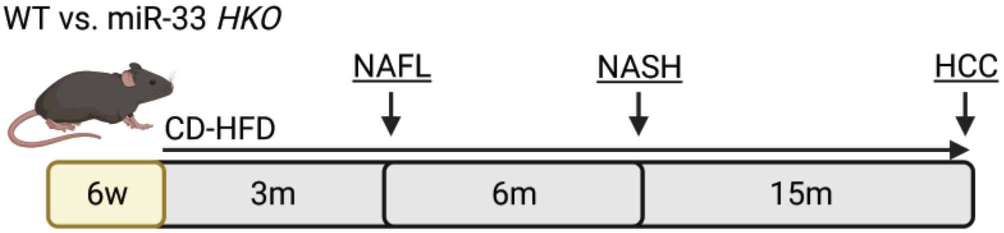


Supplemental Figure 1

Supplemental Figure 2

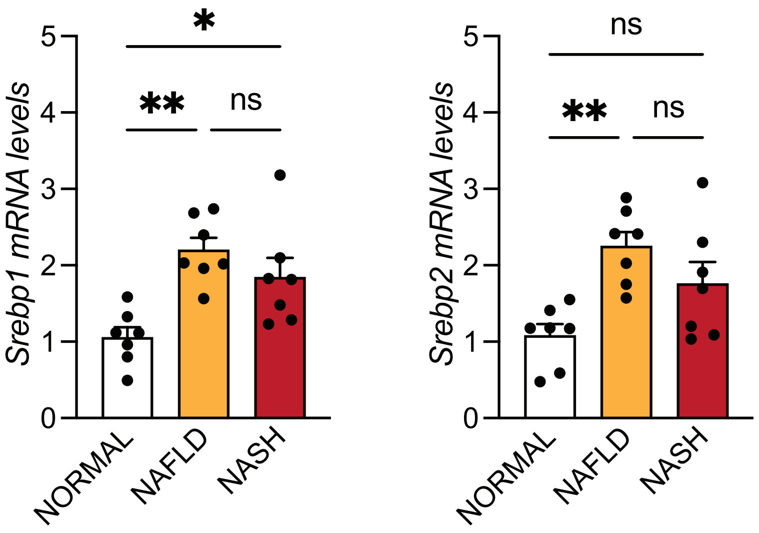

Supplemental Figure 3

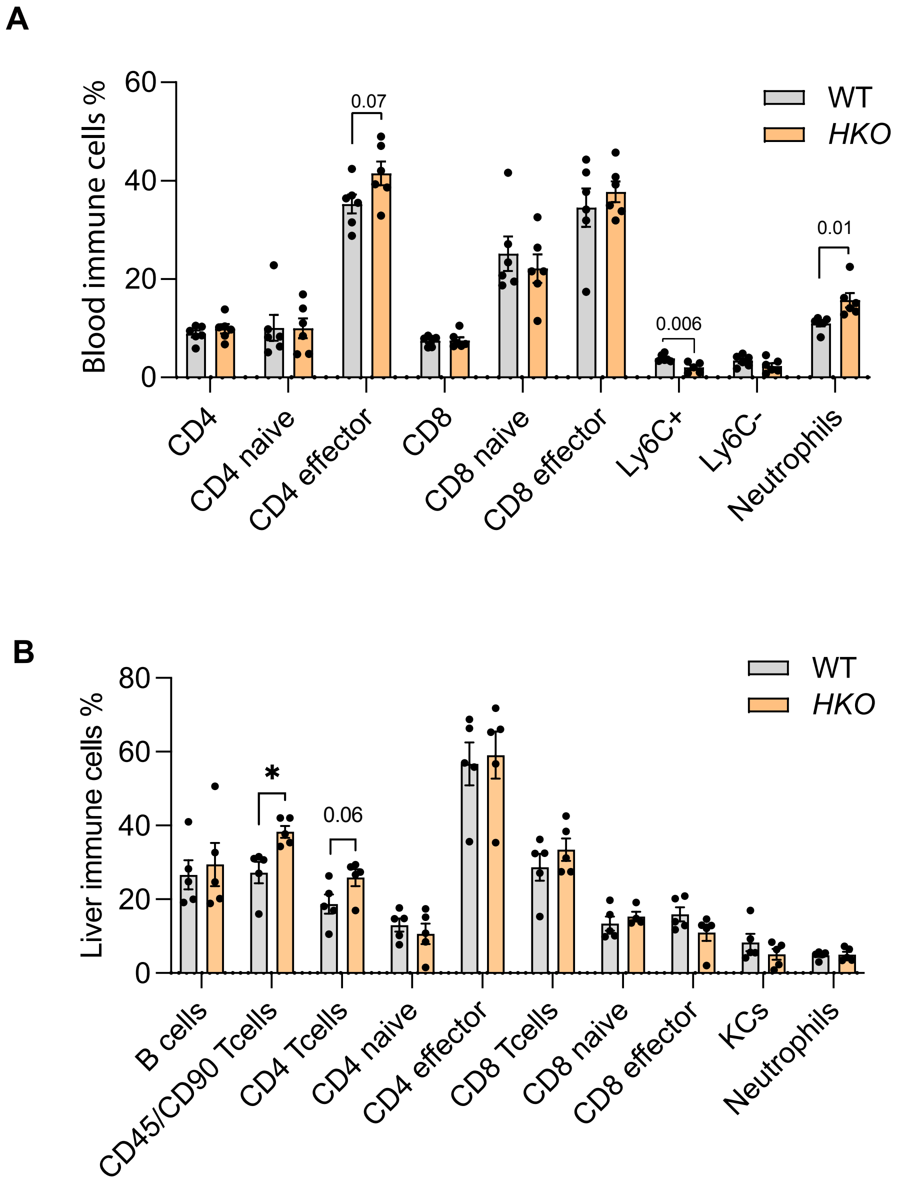

### Supplemental Figure 4

**A**

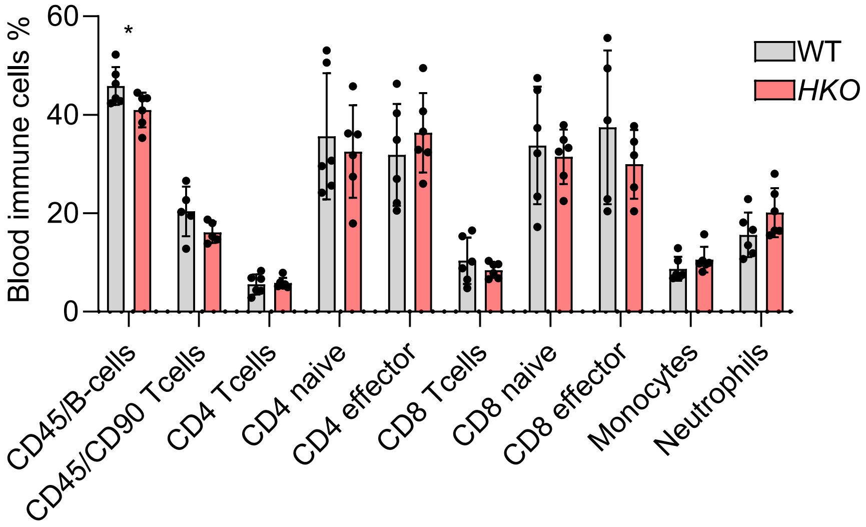


**B**

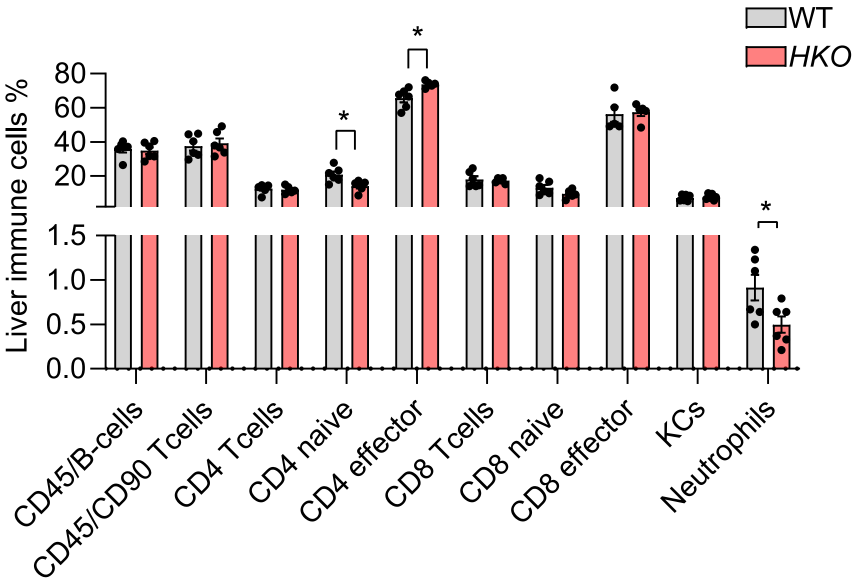

Supplemental Figure 5

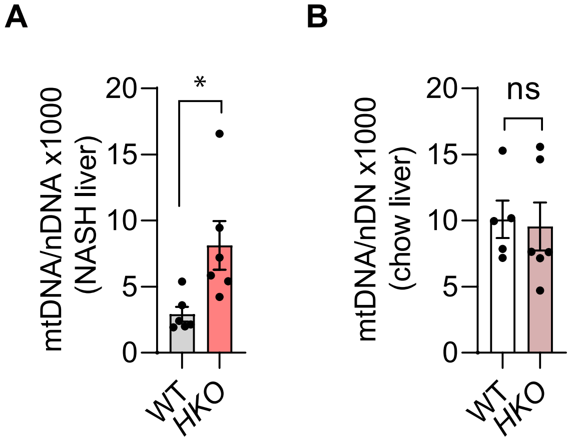

### Supplemental Figure 6

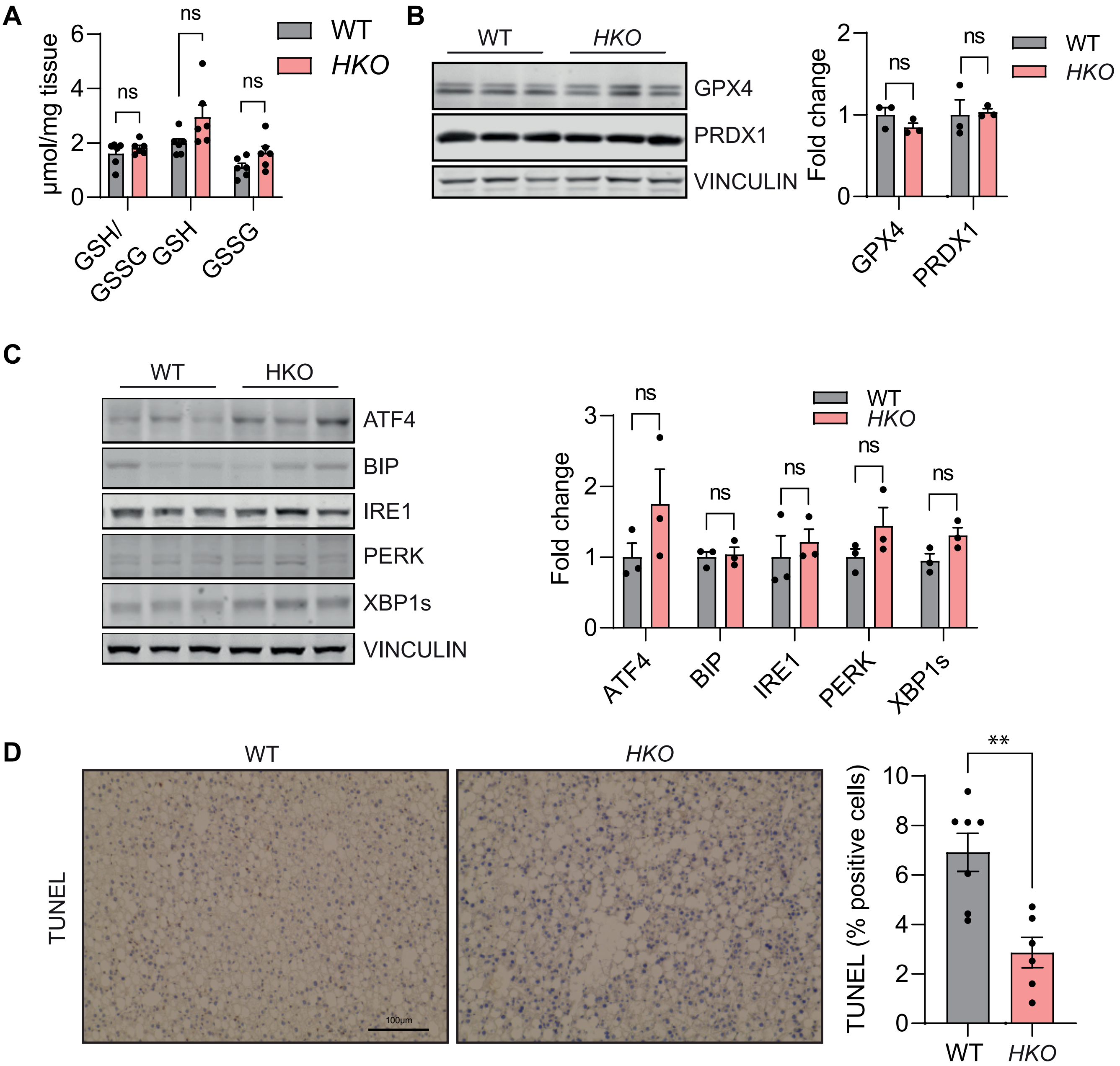

### Supplemental Figure 7

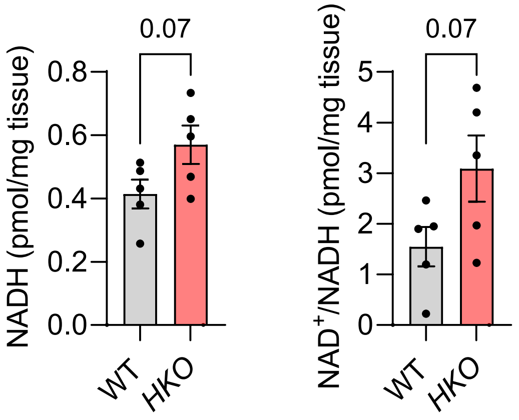
